# Heterobifunctional proteomimetic polymers for targeted protein degradation

**DOI:** 10.1101/2025.03.07.641543

**Authors:** Max M. Wang, Mihai I. Truica, Brayley S. Gattis, Julia Oktawiec, Vinay Sagar, Ananya A. Basu, Paul A. Bertin, Xiaoyu Zhang, Sarki A. Abdulkadir, Nathan C. Gianneschi

**Affiliations:** Department of Chemistry, Northwestern University, Evanston, IL, USA; Department of Biomedical Engineering, Northwestern University, Evanston, IL, USA; International Institute for Nanotechnology, Northwestern University, Evanston, IL, USA; Chemistry of Life Processes Institute, Northwestern University, Evanston, IL, USA; Department of Urology, Northwestern University Feinberg School of Medicine, Chicago, IL, USA; Robert H. Lurie Comprehensive Cancer Center, Northwestern University Feinberg School of Medicine, Chicago, IL, USA; Grove Biopharma, Inc., 1375 W. Fulton St., Ste. 650, Chicago, IL, 60558, USA

## Abstract

The burgeoning field of targeted protein degradation (TPD) has opened new avenues for modulating the activity of previously undruggable proteins of interest. To date, TPD has been dominated by small molecules containing separate linked domains for protein engagement and recruitment of cellular degradation machinery. The process of identifying active compounds has required tedious optimization and has been successful largely against a limited set of targets with well-defined, suitable docking pockets. Here we present a polymer chemistry approach termed the HYbrid DegRAding Copolymer (HYDRAC) to overcome standing challenges associated with the development of TPD. These copolymers densely display either peptide-based or small molecule-derived degradation inducers and target-binding peptide sequences for the selective degradation of disease-associated proteins. HYDRACs are synthesized in a facile manner, are modular in design, and are highly selective. Using the intrinsically disordered transcription factor MYC as an initial proof-of-concept, difficult to drug protein target, HYDRACs containing a MYC-inhibitory peptide copolymerized with a validated degron, showed robust and selective degradation of the target protein. Treatment of tumor-bearing mice with MYC-targeted HYDRACs showed decreased cell proliferation and increased tumor apoptosis, leading to significantly suppressed tumor growth *in vivo*. The versatility of the platform was demonstrated by substituting the degron for recruiters of three different E3 ligases (VHL, KEAP1, and CRBN), which all maintained MYC degradation. To demonstrate generalizability, HYDRACs were further designed against a second elusive target of clinical interest, KRAS, by employing a consensus RAS binding motif. RAS-targeted HYDRACs showed degradation in two cell lines harboring separate KRAS alleles, suggesting potential pan-KRAS activity. We envision the HYDRAC platform as a generalizable approach to developing degraders of proteins of interest, greatly expanding the therapeutic armamentarium for TPD.

## Introduction

Molecular agents which act by inducing proximity between two or more proteins have emerged as a disruptive paradigm for pharmaceutical intervention against traditionally undruggable targets ^1, 2^. The most well-known example of these compounds are inducers of targeted protein degradation (TPD), classified mainly as PROTACs (PROteolysis TArgeting Chimeras) ^3^ or molecular glues ^4^, with new, related modalities co-opting various cellular systems including autophagy or the endo-lysosomal pathway in development ^5^. The ability of these heterobifunctional entities to bring together enzymes and target proteins as neo-substrates has made them not only an exciting new therapeutic modality, but also powerful probes for chemical biology ^6^. However, the small molecule chassis and synthetic approaches around which these compounds are commonly built, limits their compositions to mostly two, or up to three ^7^, distinct domains against targets with suitable docking sites. The ability to rationally design constructs with predictable activity has also remained elusive ^8^. We hypothesized that employing copolymers, which enable multivalent display of targeting ligands and degrons covalently attached at a high repeat density, would result in a generalizable and efficient approach to TPD, stemming from a combination of ease of synthesis, cooperative binding, the direct use of targeting peptides and avidity.

To test this concept, we employed a novel proteomimetic platform technology based on peptide brush copolymers, designed to carry both targeting warheads and recruiters of cellular protein degradation machinery. We term these HYbrid DegRAding Copolymers (HYDRAC) in reference to their ability to multiplex different protein binders and degradation inducing sequences, building upon lessons learned from studies on small molecule PROTACs ^8, 9^. Peptides arranged as HYDRACs exhibit favorable emergent properties including increased resistance to proteolysis and high levels of cellular uptake as intrinsic properties of the material ^10-12^. Outlined here, we demonstrate that these attributes, paired with the modular nature of living polymerization reactions, makes the HYDRAC platform well suited for the development of targeted degraders.

We selected the proto-oncogene c-MYC (MYC), a well-validated and high value cancer target ^13, 14^, as an initial focus for the proof-of-concept studies presented here. The MYC protein is intrinsically disordered and lacks enzymatic activity, two main reasons why traditional structure-based small molecule approaches to MYC inhibition have remained elusive. To overcome this, we employed a validated MYC-inhibitory peptide ^15-17^ (H1: NELKRAFAALRDQI), derived from MYC’s first helix of the basic region/helix-loop-helix/leucine zipper (bHLH-LZ) ^18^ domain shown to interact with a binding hotspot (MYCHot) ^19^, as our targeting warhead incorporated via graft-through polymerization. Paired with this, we copolymerized recruiters of cellular protein degradation machinery, starting with a small tetrapeptide degron (RRRG) which enables rapid proteasome-mediated degradation when fused to proteins of interest ^20-23^. Substituting the degron for other recruiters of known E3 ligases (VHL, KEAP1, and CRBN) exhibited MYC degradation in H1-containing HYDRACs, validating the generalizability of the platform technology. We report here the first demonstration of a polymer capable of targeted inhibition and directed degradation of the difficult to drug protein MYC via recruitment of four different degrons, as a proof-of-concept case study for the modular HYDRAC platform with selective on-target activity and *in vivo* efficacy. Indeed, we propose this copolymer approach as a versatile and generalizable platform for rapidly converging on selective binders and active degraders of proteins of interest.

## Results

### HYDRACs containing MYC inhibitory peptide H1 stably bind MYC

To demonstrate the feasibility of HYDRACs for the selective degradation of a therapeutically challenging target, we identified the intrinsically disordered cancer-associated transcription factor MYC ^24^ as a promising test case where traditional pharmacologic attempts to modulate the target have had limited success ^25-27^. In addition to attempts at developing small molecule MYC inhibitors ^15, 28, 29^, a peptide-based binder derived from helix 1 of the bHLH-LZ region of MYC (H1) has been identified ^18^ and utilized in conjunction with different delivery strategies to improve its therapeutic potential ^30-34^. Using H1 as the targeting warhead, we designed HYDRACs containing a previously validated tetrapeptide RRRG degron sequence shown to direct tagged proteins for proteasome-mediated degradation ^35-37^ (Fig. 1a). Polymers were synthesized via graft-through ring-opening metathesis polymerization of norbornenyl-peptide monomers ^10, 38-41^ (Nor-H1, Nor-RRRG, and a scrambled version of Nor-H1, Nor-sH1 – see Supplementary Table 1), resulting in brush polymer structures consisting of a polymer backbone adorned in a stochastic manner with MYC- and proteosome-targeting sidechains (Supplementary Figs. 1-3). In this manner, a set of polymers (**P-**) were prepared, which comprised homopolymers of Nor-H1 (**P-H**), Nor-RRRG (**P-R**), and Nor-sH1 (**P-sH**), as well as copolymers of Nor-sH1 with Nor-RRRG (**P-sHR**) and our active copolymer consisting of Nor-H1 and Nor-RRRG (**HYDRAC**), targeting degrees of polymerization (DP) of 6-7 for homopolymers and 12-15 for copolymers (Fig.1a, Supplementary Table 2). In aqueous environments, these polymers fold into globular morphologies reminiscent of native proteins, driven by the hydrophobic nature of the backbone (Fig. 1b). A beneficial emergent property of this assembly is the ability to resist proteolytic degradation due to spatial hindrance of enzyme sites ^10, 42^. Using chymotrypsin as a model enzyme with a single cleavage site present in the H1 sequence, we tracked enzyme activity against the polymers using HPLC (Supplementary Fig. 4). Notably, DP influenced proteolysis with polymers containing a higher number of side chains being more resistant, independent of the specific sequence of side chains (Fig. 1c,d, P-H_long_: DP = 12, see Supplementary Table 3).

**Fig. 1.**
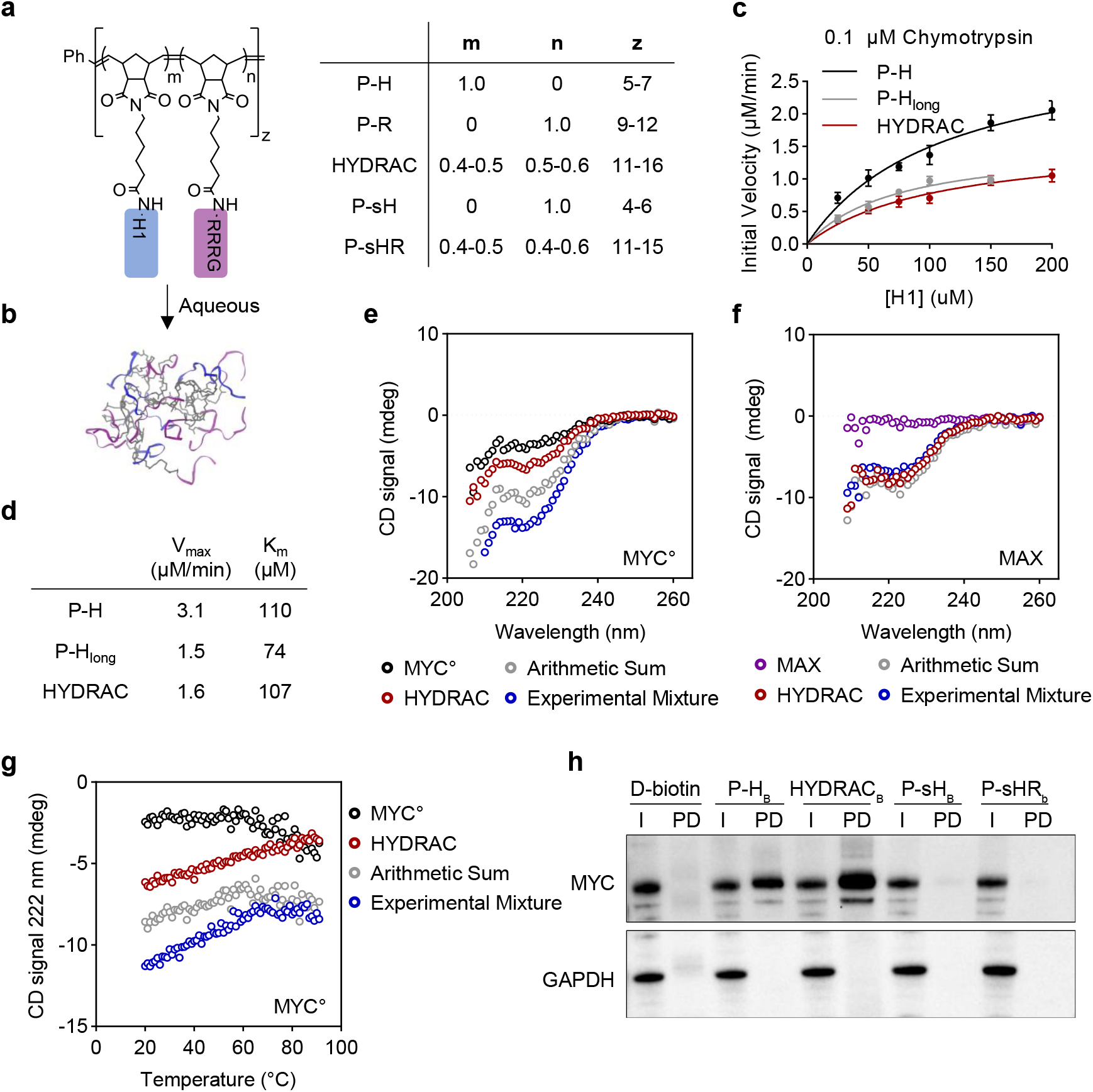
Design and validation of MYC-targeted HYDRACs. HYDRACs are heterobifunctional polymeric compounds consisting of two distinct domains, a protein-targeting ligand and a proteosome recruiting degron. The chemical nature of their synthesis allows for fine-tuned modulation of both the relative ratios and spatial distribution of each component. **a**, Chemical structure and naming convention of statistical copolymers used: P-H = H1 peptide homopolymer; P-R = RRRG degron homopolymer; HYDRAC = H1 peptide and RRRG degron copolymer; P-sH1 = scrambled H1 peptide homopolymer; P-sHR = scrambled H1 peptide and RRRG degron copolymer; m,n: proportion contribution of indicated side chain. m + n = 1; z: total polymer DP. **b**, In aqueous environments, HYDRACs collapse into a globular 3-D architecture, wherein peptide side chains are arranged around a hydrophobic polymer core. The polymerization technique yields a disperse population of structures, with each individual polymer chain having a random distribution of side chain compositions. This advantageously results in a range of possible spatial distances between any two individual side chains spanning from adjacent to the full length of the polymer. **c-d**, Michaelis-Menten plots of indicated polymer compositions in 0.1 μM chymotrypsin with cleavage rates of H1 monitored via HPLC (**c**) and calculated V_max_ and K_m_ values (**d**). P-H_long_: DP = 12 (See Supplementary Table 3 for batch characterizations). Data depict mean ± s.d. in (**c**). Experiment repeated with similar results. **e-f**, Far-ultraviolet (UV) CD spectra of the b-HLH-LZ domain of MYC (MYC°) (black, 5 μM monomer units) (**e**) or full-length Max (purple, 5 μM monomer units) (**f**), HYDRAC (red, 1 μM polymer units), the arithmetic sum of red and black spectra (gray), and the spectrum of a mixture of either protein plus HYDRAC at a 5 to 1 molar ratio, equivalent to 1 to 1 binding peptide to protein ratio (blue) recorded at 20° C. **g**, Thermal denaturation of the mixture described in (**e**). Experiment repeated with similar results. **h**, Pull down of biotin-labeled HYDRACs. PC3 cell lysates were treated with D-biotin control or indicated polymer compositions at 5 μM for HYDRAC and 10 μM for all other compositions for 2 h, after which labeled proteins underwent streptavidin pulldown, elution, separation by SDS-PAGE and blotting for MYC and GAPDH. Input lysate and pulldown lanes are shown. Representative blot from n = 3 independent experiments is shown in (**h**). X_B_: biotin-terminated polymer.

To assess MYC target engagement, we next characterized the ability of HYDRACs to heterodimerize with either the b-HLH-LZ domain of MYC (MYC°) or full length MAX using circular dichroism (CD) spectroscopy (Fig. 1e-g). The CD spectrum of the HYDRAC alone showed minima around 208 and 222 nm, characteristic of a helical structure (Fig. 1e). A pronounced gain in helical-specific signal intensity was observed only in the experimental mixture of MYC° + HYDRAC compared to the arithmetic sum of each component curve (Fig. 1e), suggesting MYC°/HYDRAC heterodimerization in the low micromolar range versus limited MAX/HYDRAC binding (Fig. 1f). Furthermore, the formed MYC°/HYDRAC heterodimers are thermodynamically stable up to 90° C (Fig. 1g). Polymers containing just the MYC-targeting H1 sequence (P-H) also showed selective heterodimerization with MYC°, while scrambled sequence controls lack the enhancement in signal indicative of binding (Supplementary Fig. 5).

Next, using biotin-terminated polymers (Supplementary Fig. 6, **P-X**_**B**_: biotin terminated polymer), we confirmed direct binding to endogenous MYC protein from cell lysates. Notably, biotin-labeled HYDRACs were able to pull down MYC protein from cell lysates at a much lower concentration compared to H1 homopolymer, P-H_B_ (Fig. 1h). This is likely due to increased electrostatic interactions between the negatively charged protein and HYDRACs which contain a polycationic RRRG degron. Polymers lacking a functional MYC-targeting sequence, both with and without the accompanying degron (P-sHR_B_, P-sH_B_) expectedly failed to stain for MYC after pull down. Taken together, results from these orthogonal methods confirm the ability of HYDRACs to engage MYC in cell-free conditions.

### MYC HYDRACS are cell penetrant and disrupt target-driven transcriptional programs

Having shown the ability of MYC-targeted HYDRACs to engage their target in cell free conditions, we next assessed cellular uptake and on-target activity *in vitro*. A Cy5.5 fluorescent labeled monomer was added to the polymers as the final monomer after complete consumption of norbornenyl-peptides to yield fluorophore labeled versions (P-H_F_, P-R_F_, P-sHR_F_, and HYDRAC_F_), which allows for tracking of internalization into cancer cells (Fig. 2a). To ensure similar fluorescence inputs between treatment groups, dosing was done based on concentration of Cy5.5 dye (Supplementary Fig. 7). MYC-sensitive PC3 prostate cancer cells were incubated with P-H_F_, P-R_F_, HYDRAC_F_, sHR_F_ at a concentration normalized to 0.5 μM with respect to Cy5.5 for 1 hour, trypsinized, and washed with heparin to remove surface bound materials ^10^. All polymer compositions tested showed high levels of uptake in PC3 cells as quantified by flow cytometry (Fig. 2b) with internalization confirmed by confocal microscopy (Fig. 2c). A combination of uptake pathways, including clathrin/caveolin-mediated endocytosis and macropinocytosis (M), are known to be critical to how both cell penetrating peptides (CPPs) and nanomaterials in general enter cells ^43, 44^. To elucidate the contributions of each of these mechanisms on the uptake of HYDRACs, PC3 and A549 cells were preincubated with a panel of endocytosis inhibitors prior to treatment with HYDRAC_F_. Incubation of both cell lines at 4 °C greatly attenuated the Cy5.5 signal, implying that HYDRACs can enter through adenosine triphosphate–dependent uptake mechanisms (Fig. 2d, Supplementary Fig. 8). FACS analysis showed slight variations in the relative importance of pathways between cell lines, with caveolin- and clathrin-mediated endocytosis playing important roles in HYDRAC uptake with lesser involvement of macropinocytosis (Fig. 2d).

**Fig. 2.**
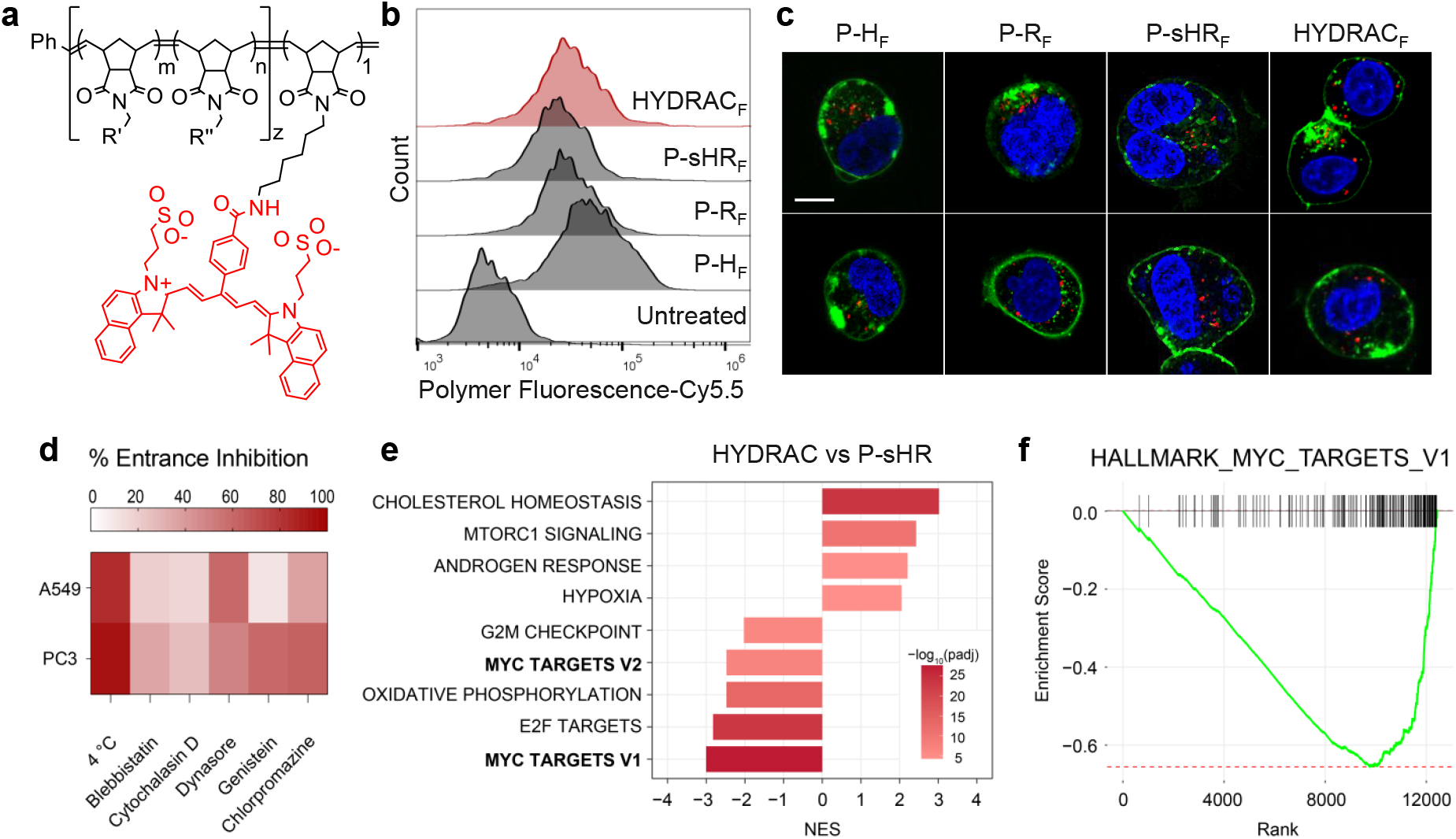
MYC HYDRACs efficiently penetrate human cancer cell lines with targeted disruption of transcriptional regulation. **a**, A Cy5.5-containing monomer (red) was attached to polymer backbone by copolymerizing 1 equivalence of dye-monomer added at the end of the reaction. **b**, Prostate cancer PC3 cells were treated with 0.5 μM (Cy5.5 conc.) of indicated polymer compositions for 1 h in media and uptake quantified by flow cytometry. Representative histogram from n = 3 independent experiments shown. **c**, PC3 cells were treated with 0.5 μM (Cy5.5 conc.) for 1 h stained with WGA-488 and Hoechst 33342, fixed, and imaged by confocal microscopy. Representative images shown. Red: Polymer-Cy5.5. Blue: Hoechst 33342. Green: WGA-488. Scale bar: 10 μm. **d**, Pharmacological and thermal probes of uptake pathways. PC3 or A549 cells were pretreated with inhibitors of macropinocytosis (100 μM blebbistatin, 10 μM cytochalasin D), clathrin-mediated (10 μM chlorpromazine, 160 μM dynasore), or caveolin-mediated endocytosis (200 μM genistein) for 15 min followed by treatment with 0.5 μM of HYDRAC-Cy5.5 for 15 min in the presence of the inhibitor, trypsinized, and analyzed by flow cytometry. Percent entrance inhibition was calculated by comparing with vehicle-treated control cells. All cells kept at 37°C unless otherwise noted. Heatmap shows summary data from n = 3 independent experiments. **e-f**, GSEA comparing gene expression profiles of scrambled H1 peptide and RRRG degron copolymer (sHR) versus HYDRAC-treated PC3 cells. Normalized enrichment scores (NES) and p values of top gene sets are listed in (**e**). Representative plot of Hallmark MYC Targets V1 pathway enrichment in HYDRAC vs sHR treated cells shown in (**f**). Experiment repeated with similar results.

We also performed transcriptome sequencing (RNA-Seq) in PC3 cells treated for 24 hours with HYDRACs, comparing transcription-level changes with P-sHR or vehicle treated cells. Transcriptional profiling of the HYDRAC treatment group compared to non-targeted scramble sequence controls showed MYC-driven gene signatures to be the most significantly downregulated pathways by gene set enrichment analysis (GSEA), suggesting HYDRAC treatment inhibits MYC-mediated transcriptional activation in cells (Fig. 2e,f). Alternatively, cells treated with non-targeting scramble controls and RRRG degron homopolymers were undifferentiable from vehicle treatment on GSEA analysis, with no significantly enriched pathways between groups (Supplementary Fig. 9). The combination of cell uptake and transcriptome results indicate that HYDRACs efficiently enter cells in a payload-independent manner primarily via active transport and subsequently exert on-target activity.

### Cellular toxicity is dependent on both side chain composition and presence of functional MYC protein

Employing three independent cancer cell lines, we assessed toxicity *in vitro* following treatment with either MYC-targeted HYRACs or scrambled controls wherein the degron is not targeted (Fig. 3a-c). All cell lines tested showed a marked difference in susceptibility to HYDRACs compared to P-sHR with a near order of magnitude difference in IC_50_ separating the two treatments. To differentiate the contributory effects of each HYDRAC component on the observed toxicity, a series of polymers were synthesized and toxicity assessed in two independent cell lines. HYDRAC treatment consistently showed significantly lower IC_50_ values compared to P-H, P-R, or the combination treatment of the two, with P-sH and P-sHR showing almost no toxicity, highlighting the importance of both domains needing to be conjugated to the same polymer backbone (Fig. 3d). Notably, P-H had a much lower IC_50_ value (∼10 μM) compared to unpolymerized free H1 peptide (∼700 μM), likely resulting from avidity effects and improved cell uptake when assembled as a polymer. P-R (synthesized at slightly higher DPs given the small monomer size and difficulty in reliably making lower molecular weight polymers, see Supplementary Table 3) itself displayed some cellular toxicity being attributed to off-target effect of the RRRG degron itself (Fig. 3d). Encouragingly, we observed single digit micromolar IC_50_ values exclusively in HYDRAC treated cells, in alignment with the dose range required for robust effects (within 24 h) on MYC transcriptional program (Fig. 2e-f). HYDRAC treated PC3 cells also showed higher levels of apoptosis compared to P-sHR, as assessed by Annexin V/PI staining (Fig. 3e), adding credence to the evidence that cancer cells are susceptible to HYDRAC therapy.

**Fig. 3.**
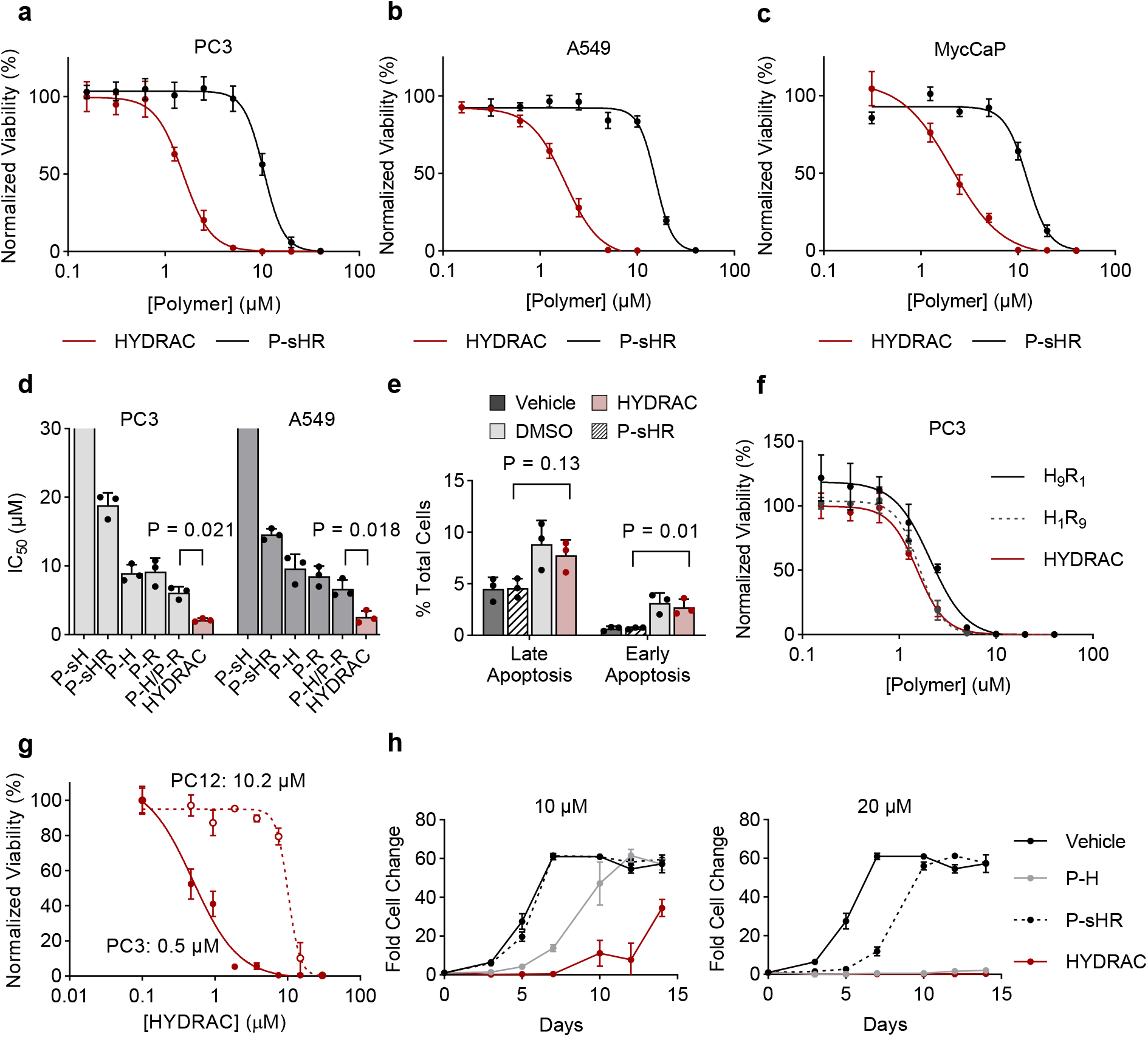
Toxicity following HYDRAC treatment is formulation- and MYC-dependent. **a-c**, Differential cell toxicity following treatment with HYDRAC or P-sHR. Representative dose-response plots of PC3 (**a**), A549 (**b**), or MycCaP (**c**) cells following 72 h treatment with viability normalized to vehicle treated control quantified by CTGlo. **d**, Summary IC_50_ values in PC3 and A549 cells following treatment with indicated polymers averaged from n = 3 independent biological experiments with > 2 different batch formulations. sH treated cells had IC_50_ values >40 μM. **e**, Annexin V/PI staining of PC3 cells treated for 24 h with HYDRAC analyzed by flow cytometry and split into late (Annexin V-FITC ^+^/PI ^+^) and early (Annexin V-FITC ^+^/PI ^+^) stage apoptosis. DMSO group was incubated with 10% DMSO as a positive control. n = 3 independent samples per group. Experiment repeated with similar results. **f**, Representative dose response curves of PC3 cells treated for 72 h with HYDRACs consisting of different targeting to degron ratios with viability normalized to vehicle treated controls. **g**, PC3 (solid) or PC12 (dashed) cells were treated with HYDRAC for 72 h and cell viability normalized to vehicle treatment. IC_50_ values listed. **h**, A549 cells were incubated with a single treatment of 10 or 20 μM of indicated polymer formulations at day 0 and cell counts normalized to initial seeding amount monitored over 2 weeks. Data depict mean ± s.d. P-values determined by one-way ANOVA in **d, e**. One representative dose response or proliferation curve from n = 3 independent experiments shown in **f, g**, and **h**.

Having narrowed down on a lead HYDRAC composition consisting of equimolar ratios of H1 to RRRG, we next explored if varying the targeting warhead to degron ratio improved efficacy. PC3 cells treated with HYDRACs (1H1: 1RRRG) were compared to cells treated with polymers containing either a 1:9 (H_1_R_9_) or 9:1 (H_9_R_1_) H1:RRRG ratio. All tested ratios showed similar levels of cellular toxicity (Fig. 3f), suggesting that the optimal spacing between the two domains is encompassed within the relative spacings of the random copolymer. To confirm observed toxicity effects were on-target, we compared effects of HYDRAC treatment on a PC12 cell line which lacks the MYC dimerization partner MAX ^45^. In stark contrast to PC3 cells, PC12 cells resisted treatment with MYC-targeted HYDRACs, supporting the conclusion that observed cellular toxicities are MYC-dependent (Fig. 3g).

With HYDRACs displaying resistance to enzymatic degradation in a model enzyme system (Fig. 1c), we aimed to determine if these compounds display long-acting effects *in vitro*. To assess this, A549 cells were incubated with a single treatment of either 10 or 20 μM HYDRAC or control polymers and tracked over two weeks. Both P-H and HYDRAC treatments suppressed cell growth in the short term, with HYDRAC treated cell counts not recovering until 2 weeks post treatment, and then at a much slower growth rate (Fig. 3h). This correlates well with trends observed in both the enzymatic degradation and cellular toxicity studies, likely a result of the combination of HYDRACs being both longer circulating and more potent compared to P-H. P-sHR, meanwhile, only mildly retarded cell proliferation at a single treatment of 20 μM (Fig. 3h), consistent with the lower observed IC_50_ values (Fig. 3d). Satisfied with these results demonstrating HYDRACs induced on-target effects on cell growth, we next explored the ability of these constructs to selectively degrade the target protein.

### HYDRACs selectively degrade their target in a catalytic manner

P-H (Nor-H1 homopolymers lacking the RRRG degron) failed to reduce endogenous MYC levels at doses up to 20 µM, at which point cellular toxicity effects are evident (Fig. 4a). In stark contrast, MYC-targeted HYDRACs exhibit noticeably reduced MYC protein bands by Western Blot at doses as low as 0.6 μM, with non-targeting degron containing polymers (P-sHR) showing no protein level reductions, even at ∼20X the dose (Fig. 4a). These observed changes in MYC protein levels are not caused by changes in MYC mRNA levels, which remained unchanged following HYDRAC treatment (Supplementary Fig. 10). Treatment of HEK293T cells with HYDRACs displayed similar dose-dependent MYC degradation, with near total ablation of MYC protein bands at high concentrations, confirming activity was cell-line agnostic (Supplementary Fig. 11). Neither treatment with either domain alone (P-H, P-R), combination treatment with homopolymers of both domains unlinked (P-H + P-R), or scrambled targeting sequence control (P-sH-R) showed appreciable MYC degradation, highlighting the importance of linking both domains on a HYDRAC for activity (Fig. 4b, Supplementary Fig. 12).

**Fig. 4.**
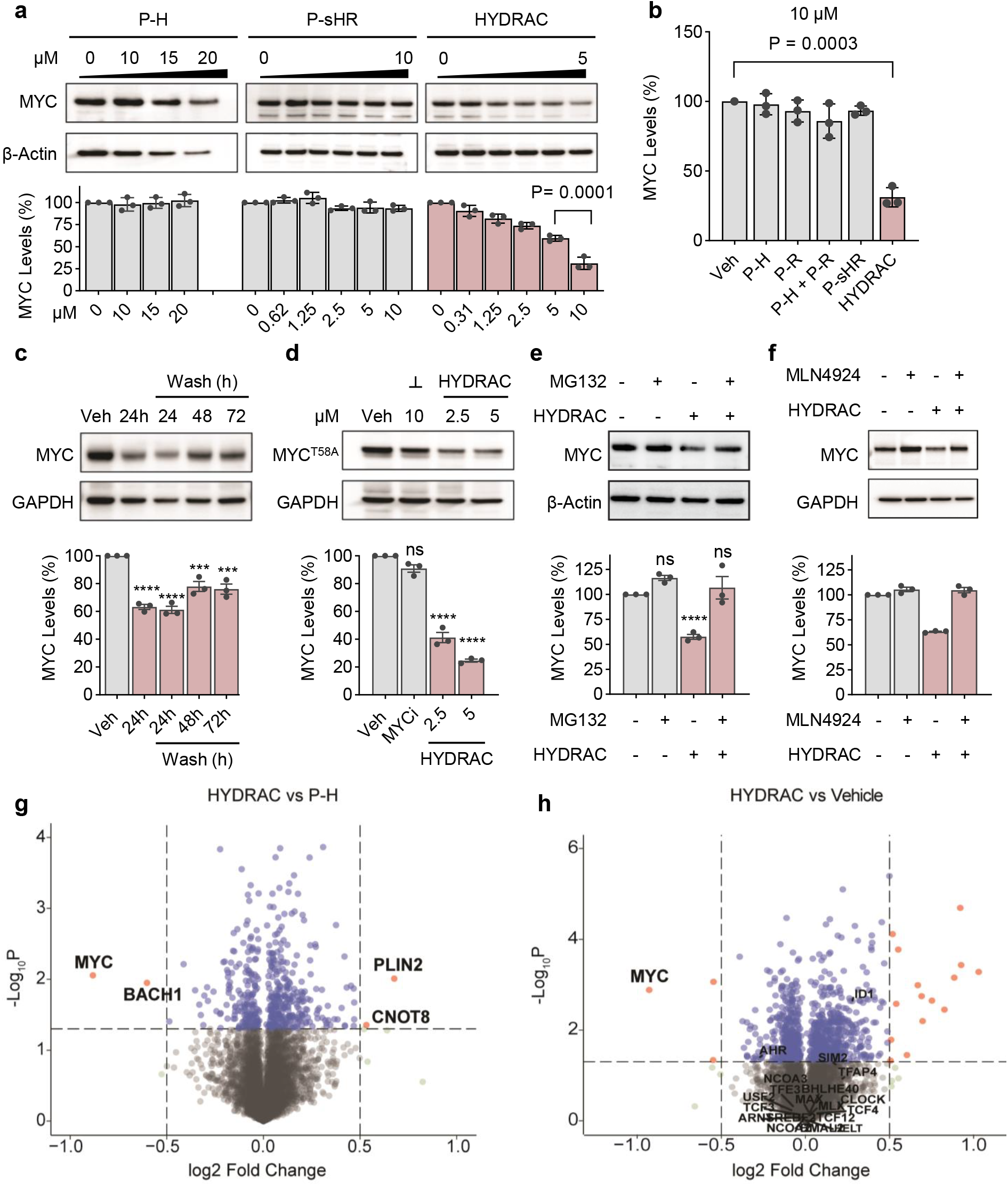
Characterizing the mechanism of HYDRAC induced MYC degradation. **a**, Representative western blot and quantification of endogenous MYC protein levels in PC3 cells treated for 24 h with increasing doses of P-H, P-sHR, or HYDRACs. 20 µM P-H treatment resulted in significant levels of observed cell death. **b**, An intact HYDRAC containing targeting warheads conjugated with degrons is required for protein degradation. Quantification of endogenous MYC levels in PC3 cells following 24 h treatment with indicated polymer formulations at 10 µM. P-H + P-R: mixture of homopolymers. **c**, HYDRACs exhibit a catalytic mode of action. PC3 cells treated with 10 µM HYDRACs for 24 h followed by a washout and monitoring for up to 72 h were compared to continuous incubation with 10 µM HYDRAC (24 h) and vehicle treated controls. **d-f**, HYDRAC-induced MYC degradation bypasses endogenous degradation pathways and is both proteasome- and neddylation-dependent. Exogenous MYC ^T58A^ levels constitutively expressed in stably transfected PC3 cells following treatment with indicated concentrations of HYDRACs or small molecule MYC inhibitor MYCi975 (***┴***). Endogenous WT MYC visible as faint band below MYC ^T58A^ (**d**). PC3 cells were co-treated with 5 µM HYDRAC and 1 µM proteasome inhibitor MG132 (**e**) or neddylation inhibitor MLN4924 (**f**) and rescue of MYC protein levels assessed after 24 h (MG132) or 4 h (MLN4924). **g-h**, TMT-based whole proteosome quantification of HYDRAC treatment. PC3 cells were treated with vehicle, 5 µM HYDRAC, or 5 µM P-H for 24 h. Comparisons were made between HYDRACS vs P-H (**g**) and HYDRAC vs vehicle (**h**). Blots in **a**,**c-f** are representative from n = 3 independent experiments. Data in **a-f** depict mean ± s.d. from n = 3 biologically independent samples per group. P-values determined by one-way ANOVA and compared with vehicle controls unless otherwise indicated. ***: P < 0.001, ****: P < 0.0001, ns: not significant.

Inspired by the numerous reports on PROTAC formulations showing catalytic properties ^46-48^, we endeavored to test if HYDRACs act in a similar fashion. To test this, cells were treated for 24 h with HYDRACs or a vehicle control, prior to being rinsed with PBS to washout extracellular compound and media replaced with fresh polymer-free media, followed by monitoring of MYC protein levels at 24 hour intervals, for up to a total of 72 hours. HYDRAC treated cells showed sustained MYC suppression up to 72 h post washout (Fig. 4c). This observation could partially be attributed to the stability of HYDRACs, but is also consistent with a catalytic mode of action, as the rapid turnover of MYC would otherwise be expected to quickly exhaust the small amount of internalized HYDRAC ^49^.

Endogenous MYC is partially regulated through phosphorylation at threonine 58 (pT58), primed by a phosphorylation cascade initiated by several kinases, ultimately leading to E3 ubiquitin ligase mediated degradation ^50, 51^. Using a MYC ^T58A^ mutant cell line, we examined if HYDRAC treatment alters MYC stability through this mechanism. As a comparative control, we included a small-molecule MYC inhibitor (MYCi) ^15^ known to induce MYC degradation in a pT58-dependent manner. PC3 MYC ^T58A^ cells treated for 6 h with both 2.5 and 5 μM HYDRACs showed significant decreases in MYC ^T58A^ levels, evidence that HYDRAC activity acts through non-endogenous degradation pathways. MYCi treatment meanwhile showed negligible changes in mutant MYC levels, consistent with its reported mechanism of action (Fig. 4d).

As the mechanism by which the RRRG sequence used in these initial formulations acts as a degron remains poorly understood ^23, 35, 36^, we treated PC3 cells with HYDRACs in the presence or absence of proteosome and neddylation inhibitors (MG132 or MLN4924 respectively) to elucidate pathways involved in the observed protein degradation. Pretreatment with either inhibitor rescued MYC protein levels, suggesting HYDRAC activity is dependent on both the proteasome and Cullin-RING ubiquitin ligases (Fig. 4e,f).

### Whole proteome analyses show HYDRAC-induced protein changes are selectively on-target

To validate the observed HYDRAC-induced MYC degradation on Western blot and assess proteome-wide perturbations in an unbiased manner, we performed a tandem mass tag (TMT)-based quantitative proteomic analysis on HYDRAC treated PC3 cells ^52^. MYC levels were identified as the most decreased hit by mass spec in HYDRAC vs P-H treated cells, with otherwise minimal changes in protein profiles (Fig. 4g). Outside of MYC, no other BHLH protein family members (including MAX) showed significant changes both versus P-H (Supplementary Fig. 13a) or versus vehicle (Fig. 4h, Supplementary Fig. 13b). Furthermore, TMT analysis of P-H treated cells compared to vehicle failed to show downregulation of MYC protein, in line with our Western Blot results and confirming that the P-H without the RRRG degron does not degrade MYC (Supplementary Fig. 13c). Both HYDRAC and P-H treatments resulted in similar sets of significantly upregulated proteins compared to vehicle treatment, suggesting those hits arise from MYC inhibition (Supplementary Fig. 13b,c). Encouraged by the clean proteomic profile showing that the observed HYDRAC effects were indeed on-target, we next moved to test activity in a MYC-driven tumor model *in vivo*.

### MYC HYDRACs inhibit tumor growth in vivo

Having demonstrated the ability of MYC-targeted HYDRACs to selectively act *in vitro*, we explored their activity in a MYC-responsive tumor model (Fig. 5). For this, we assessed efficacy in mice bearing established MycCap tumors, previously validated to be sensitive to HYDRAC treatment (Fig. 3c) as an immunocompetent tumor model ^15^. Treatment with 25 mg/kg HYDRACs given IP three times a week (Fig. 5a) resulted in significantly suppressed tumor growth (Fig. 5b) with no detrimental changes in body weight between treatment arms (Supplementary Fig. 14). Immunofluorescent (IF) staining and immunohistochemistry (IHC) of tumors excised at day 25 post implantation showed significant reduction in proliferative cells, assessed by Ki67 staining, with high levels of cleaved caspase-3 in HYDRAC treated animals (Fig. 5c). The significant degree of anti-proliferative effects in the tumor (tumor growth inhibition of 52% at D25) underscores the therapeutic potential of the platform technology.

**Fig. 5.**
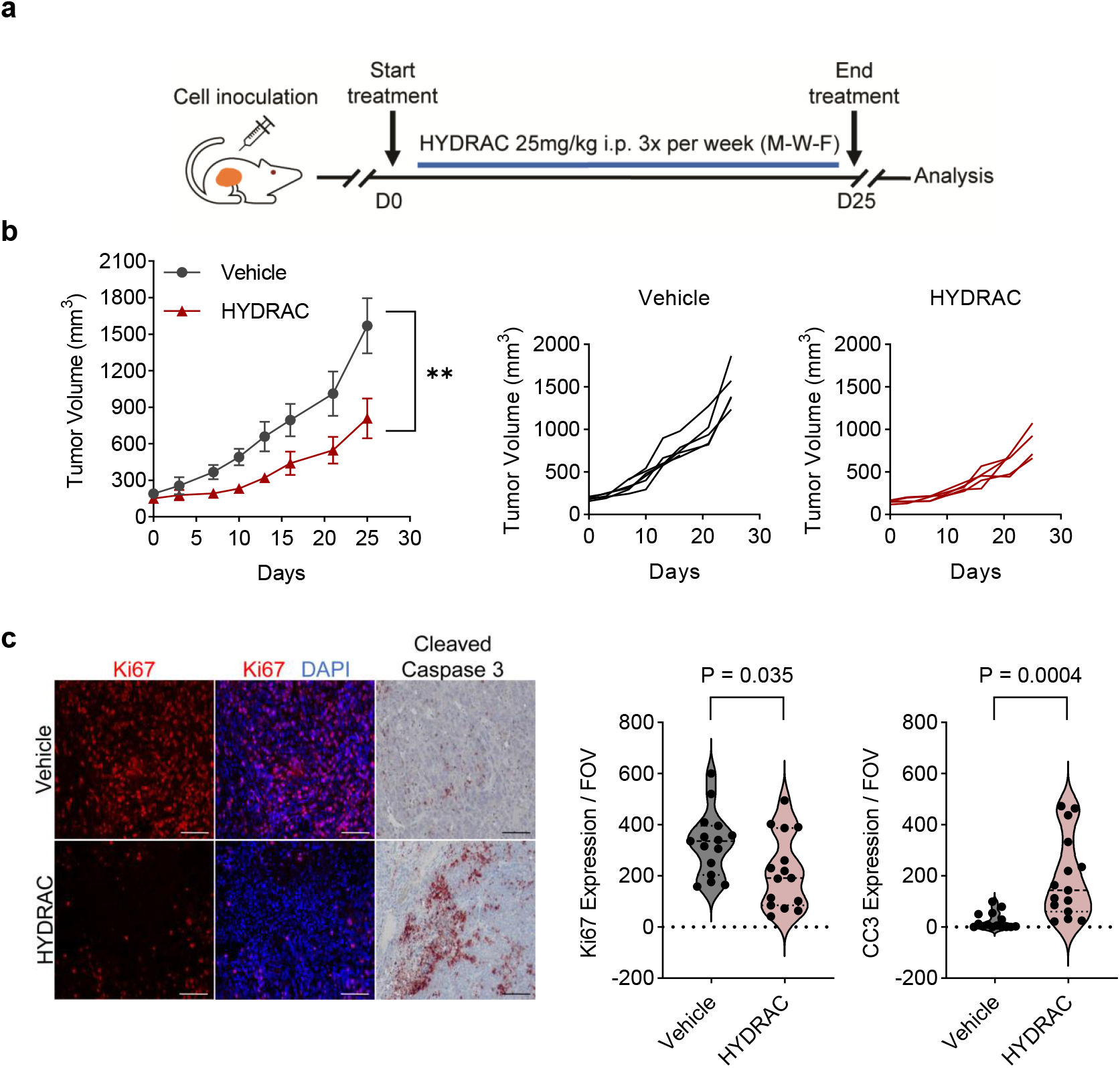
MYC-targeted HYDRACs have anti-proliferative effects in a MYC-driven tumor model. **a**, Mice bearing MycCap allografts were treated 3 times a week with vehicle or HYDRACs (25 mg/kg IP). **b-c** Average and individual tumor growth curves (**b**) and representative Ki67 IF or cleaved Caspase-3 IHC images with quantification on tumors harvested 25 days after treatment start (**c**). Scale bar: 100 um. n = 5 mice per group. Data depict mean ± s.d. P-values determined by Student’s t-test in **b**. **: P < 0.01.

### HYDRACS are capable of engaging a wide range of functional E3 ligase recruiters

One distinct advantage of the HYDRAC platform is its modular nature, being amenable to the incorporation of a wide range of side chain peptide sequences. Capitalizing on this feature, we generated a library of MYC-targeted HYDRACs switching out the RRRG degron for recruiters of different E3 ligases. We selected three validated degrons: peptide recruiters of VHL ^53^ or KEAP1 ^54^ (Supplementary Fig. 15), and the CRBN binding small molecule thalidomide ^55^ to test (Supplementary Fig. 16). With optimal MYC degradation achieved at a one-to-one targeting warhead to RRRG degron ratio, initial compositions were again synthesized at equimolar ratios of each domain (Supplementary Fig. 17). Unfortunately, thalidomide HYDRACs formed aggregates and were insoluble at equimolar ratios. Therefore, a 15:1 (HYDRAC_CRBN1_) and 10:5 (HYDRAC_CRNB2_) Nor-H1 to Nor-Thalidomide ratio were tested to maintain similar molecular weights between groups (Fig. 6a, Supplementary Fig. 18). Satisfyingly, HYDRACs containing either of the three E3 ligase recruiters all showed significant levels of MYC degradation at a concentration of 10 μM (Fig. 6b,c). Intriguingly, HYDRAC_KEAP_, HYDRAC_CRBN1_, and HYDRAC_CRBN2_ failed to decrease MYC levels at higher (20 μM) doses, suggesting potential presence of a signature “hook effect” seen commonly in PROTAC response curves wherein binding sites at protein targets and E3 ligases become increasingly saturated, preventing ternary complex formation ^56^. While further optimizations on side chain composition and mechanistic probing of degradation pathways warrant further study, these data confirm the generalizability of the HYDRAC platform and spotlight the potential “plug and play” nature of the approach.

**Fig. 6.**
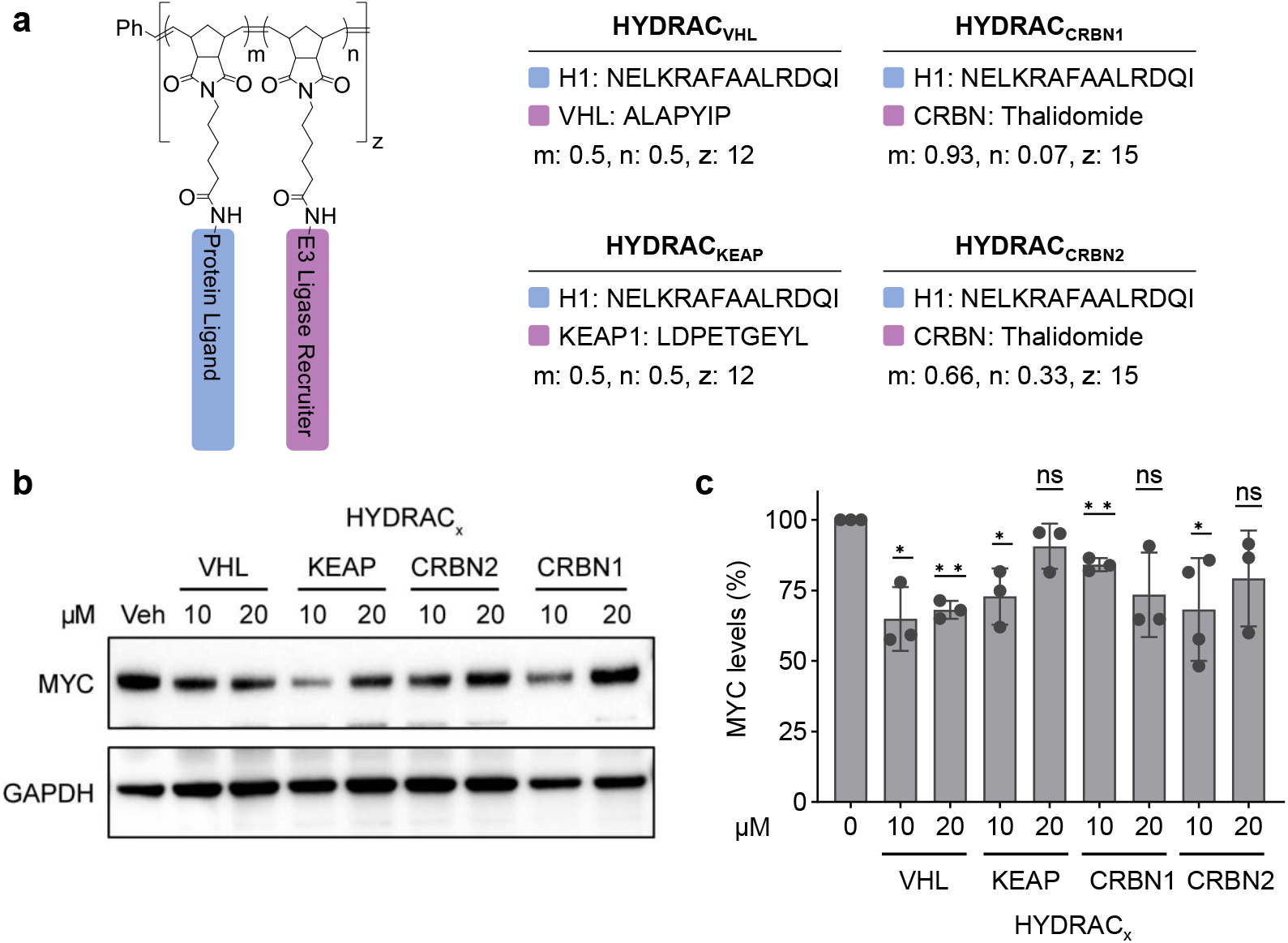
HYDRACs can recruit multiple different E3 ligases to degrade their target. **a**, Chemical structures of four MYC-targeting (blue) HYDRACs incorporating three different E3 ligase recruiting peptides or small molecule thalidomide (purple) copolymerized with H1. m,n: proportion contribution of indicated side chain. m + n = 1; z: total polymer DP. **b-c**, Representative western blot (**b**) and quantification (**c**) of endogenous MYC protein levels after treatment of PC3 cells for 24 h with indicated HYDRAC formulations and concentrations. n = 3-4 biologically independent experiments per group. Data depict mean ± s.d. P-values determined by Student’s t-test compared to vehicle-treated controls in **c**. *: P < 0.05, **: P < 0.01, ns: not significant.

### HYDRACs can be generalized to degrade a secondary target, RAS

To confirm the platform can be generalized to degrade other proteins of interest, we designed HYDRACs targeting the oncoprotein RAS using a consensus RAS binding sequence derived from RAF-1 ^57^ (Supplementary Table 1, Supplementary Fig. 19). Again, a library of HYDRACs recruiting different E3 ligases as well as RRRG (comparable to compositions targeting MYC) were synthesized. NCI-H727 cells, a bronchial carcinoid cell line carrying heterozygous KRAS G12V mutations ^58, 59^, were treated with RAS-targeted HYDRACs or controls and KRAS levels assessed by WB. Selective KRAS degradation was seen in HYDRACs containing the RRRG degron, as well as recruiters of VHL and CRBN, but not when paired with recruiters of KEAP1 (Fig. 7). We hypothesize this could be due to a loss of KEAP1 function, a common occurrence in cancer cell lines ^60^. Lastly, to assess the ability of RAS-targeted HYDRACs to act as a pan-KRAS degraders, we incubated Panc-1 cells harboring a separate KRAS G12D mutation with a library of HYDRACs (Supplementary Fig. 20). Remarkably, significant levels of KRAS degradation were also observed in this cell line, suggesting induced degradation by RAS-HYDRACs could be agnostic to mutation status. Together, these data confirm the generalizability of the HYDRAC platform and spotlight the potential “plug and play” nature of the HYDRAC platform.

**Fig. 7.**
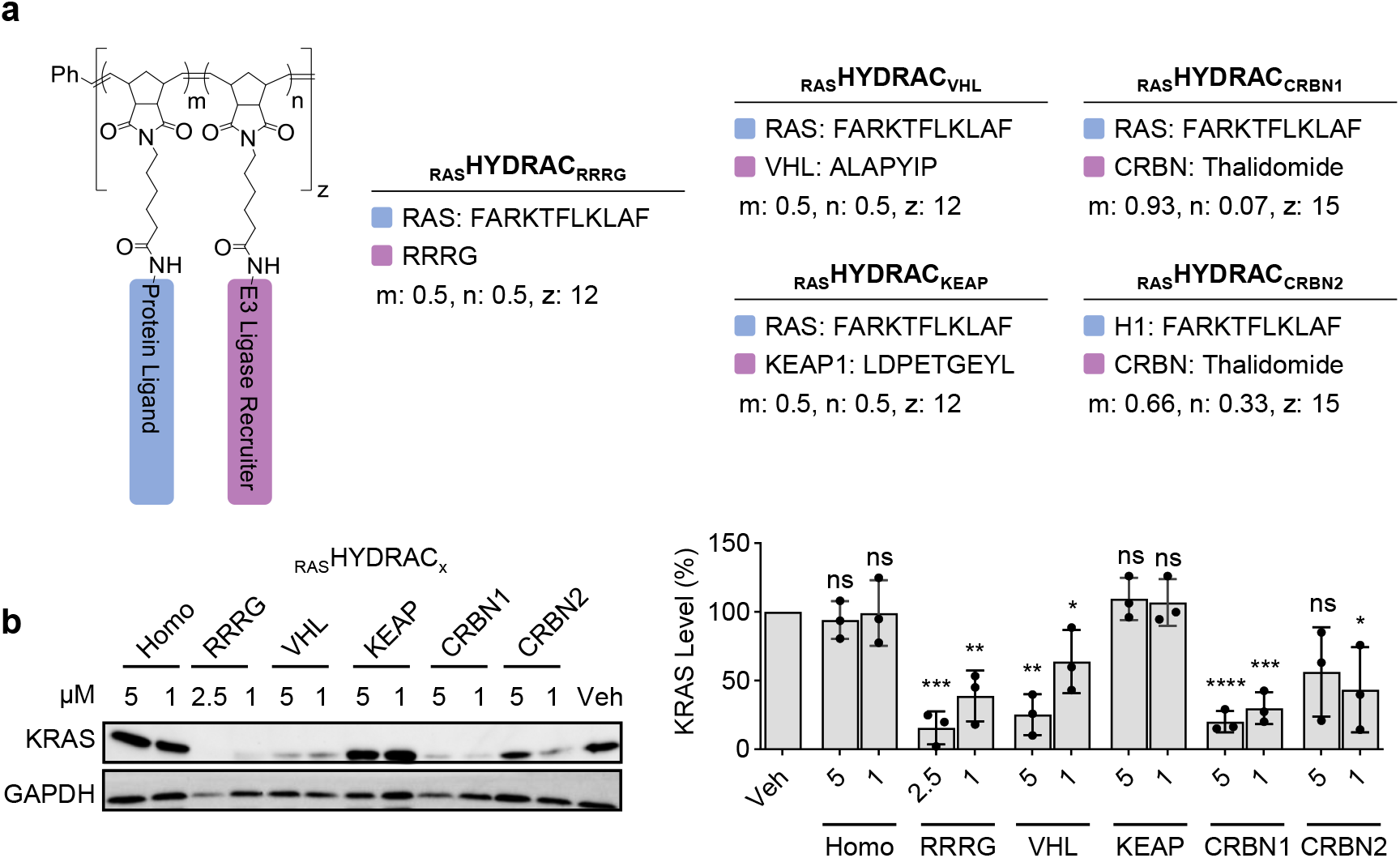
RAS targeted HYDRACs. **a**, Chemical structures of five RAS-targeting (blue) HYDRACs incorporating four different E3 ligase recruiting peptides or small molecule thalidomide (purple) copolymerized with RAS targeting peptide sequence. m,n: proportion contribution of indicated side chain. m + n = 1; z: total polymer DP. **b**, Representative western blot and quantification of endogenous KRAS protein levels after treatment of NCI-H727 cells for 24 h with indicated HYDRAC formulations and concentrations. n = 3 biologically independent experiments per group. Data depict mean ± s.d. P-values determined by Student’s t-test compared to vehicle-treated controls. *: P < 0.05, **: P < 0.01, ***: P < 0.001, ****: P < 0.0001, ns: not significant.

## Discussion

Current approaches to designing compounds for targeted protein degradation are limited by challenges inherent to small molecule chemistry, namely the need to find suitable docking sites and the inability to incorporate more than two distinct domains. Although recent efforts in the development of trivalent PROTACs ^7^ and peptide-based protein binders ^61^ have begun to address some of these challenges, there remains a need for a modular platform that can be quickly reprogrammed to target disparate disease-driving proteins. In this study, we introduce the first iteration of a class of heterobifunctional polymers, termed HYDRACs, capable of simultaneous direct inhibition and targeted degradation of proteins. Unlike currently available molecules, the polymeric nature of HYDRACs naturally results in structures encompassing a wide range of spatial distances between any two domains, spanning from directly adjacent (6 Å) to the length of the polymer chain (<15 nm). The distance between immediately adjacent sidechains is approximately 24-atoms from amide-linked peptide through the polymer backbone to the adjacent peptide (Fig. 1a), accessing potentially favorable dimensions for induced proximity effects ^62^. In addition, the three-dimensional HYDRAC globular structure exists at single-digit nanometer length scales comparable to native proteins (Fig. 1b). This, coupled with backbone flexibility and the stochastic nature of the peptide display, leads to a system capable of exploring conformational space for induced fit at binding domains on the target proteins which can self-sort to achieve optimal ternary complex formation. In stark contrast, small molecules lock the separation of separate binding and active domains in a rigid manner requiring iterative optimization of linker lengths which can be largely target-specific ^56^. In short, ongoing studies will determine design rules for optimizing HYDRAC efficacy against protein targets, with these initial results suggesting that it may be possible to bypass composition iterations as the random distribution of side chains along the polymer chassis encompasses the required relative spatial distances between domains.

In this work, we demonstrate that HYDRACs can be designed to engage challenging protein targets, leveraging the inherent multivalent display of sidechain repeats for avidity and cooperativity. As referenced above, a recent study on trivalent PROTACs with two BET inhibitors and a E3 ligase recruiter showed higher levels of degradation compared to bivalent compounds engaging the same target, which the authors attributed to increasing binding valency ^7^. Meanwhile, HYDRACs comprise multiple copies of each domain, showing selective degradation solely when targeting sequences are linked to degron motifs on the same polymer backbone. These features cannot be easily recapitulated by small molecules, where stepwise synthesis, as opposed to high efficiency polymerization, is required. To our knowledge, this is the first reported utilization of H1 as a targeting warhead to generate compounds capable of both selective inhibition and targeted degradation of MYC, a disease target which has largely eluded efforts at small molecule modulation ^19^.

The successful initial proof of concept efforts to degrade two challenging cancer associated proteins, MYC and KRAS, shown in this study opens a vast design space for other promising disease areas for this technology. The unique ability of the HYDRAC platform to rapidly swap out constituent components in a facile manner to either recruit alternative degradation pathways or target other proteins, makes it applicable to many disease states which could benefit from selective protein degradation, including non-cancer related targets such as amyloid β in Alzheimer’s disease ^63^, IRAK family members in inflammatory diseases ^64^, and HMG-CoA reductase to treat cardiovascular disease ^65^. We anticipate the HYDRAC platform will have broad applicability targeting a wide range of disease-associated proteins, constituting a new tool for chemical biology and a modality for the development of future therapeutics.

## Acknowledgements

This work was made possible by a grant from the Willens Center for Nano Oncology, International Institute of Nanotechnology (IIN) at Northwestern University and generous support from the Liz and Eric Lefkofsky Innovation Research Award. Work was performed at the Mouse Histology and Phenotyping Laboratory, which is funded by NCI CCSG P30 CA060553 awarded to the Robert H Lurie Comprehensive Cancer Center. This work also made use of NUseq, Keck Biophysics Facility, Biological Imaging Facility, the Robert H. Lurie Comprehensive Cancer Center Flow Cytometry core, and the Integrated Molecular Structure Education and Research Center (IMSERC), which receives support from the Soft and Hybrid Nanotechnology Experimental (SHyNE) Resource (NSF ECCS-2025633), IIN, and Northwestern University. M.M.W. was supported by a NIH fellowship (5F30CA257519). M.I.T. was supported by a NIH fellowship (5F30CA250196). B.S.G. was supported by a NSF fellowship (NSF-GRFP No. DGE-1842165). We would like to thank M. Thompson for assistance with peptide synthesis

## Author Contributions

M.M.W. and N.C.G. conceived the project with contributions from P.A.B. M.M.W. with M.I.T., B.S.G., and J.O. performed the experiments. A.A.B. and X.Z. performed proteomics analysis. V.S. assisted with animal studies. S.A.A. and N.C.G. supervised research and aided in designing experiments with M.I.T. and M.M.W. M.M.W. and M.I.T. produced the figures, analyzed and interpreted results, and drafted the manuscript with input from all authors. All authors discussed the results and commented on the manuscript.

## References

1. Deshaies, R.J. Multispecific drugs herald a new era of biopharmaceutical innovation. Nature 580, 329–338 (2020).

2. Stanton, B.Z., Chory, E.J. & Crabtree, G.R. Chemically induced proximity in biology and medicine. Science 359 (2018).

3. Burslem, G.M. & Crews, C.M. Proteolysis-Targeting Chimeras as Therapeutics and Tools for Biological Discovery. Cell 181, 102–114 (2020).

4. Dong, G., Ding, Y., He, S. & Sheng, C. Molecular Glues for Targeted Protein Degradation: From Serendipity to Rational Discovery. Journal of Medicinal Chemistry 64, 10606–10620 (2021).

5. Alabi, S.B. & Crews, C.M. Major advances in targeted protein degradation: PROTACs, LYTACs, and MADTACs. Journal of Biological Chemistry 296, 100647 (2021).

6. Spradlin, J.N., Zhang, E. & Nomura, D.K. Reimagining Druggability Using Chemoproteomic Platforms. Acc Chem Res 54, 1801–1813 (2021).

7. Imaide, S. et al. Trivalent PROTACs enhance protein degradation via combined avidity and cooperativity. Nature Chemical Biology 17, 1157–1167 (2021).

8. Bondeson, D.P. et al. Lessons in PROTAC Design from Selective Degradation with a Promiscuous Warhead. Cell Chemical Biology 25, 78-87.e75 (2018).

9. Gao, H., Sun, X. & Rao, Y. PROTAC Technology: Opportunities and Challenges. ACS Medicinal Chemistry Letters 11, 237–240 (2020).

10. Blum, A.P. et al. Peptides Displayed as High Density Brush Polymers Resist Proteolysis and Retain Bioactivity. Journal of the American Chemical Society 136, 15422–15437 (2014).

11. Sun, H. et al. Origin of Proteolytic Stability of Peptide-Brush Polymers as Globular Proteomimetics. ACS Central Science 7, 2063–2072 (2021).

12. Choi, W. et al. Biomolecular Densely Grafted Brush Polymers: Oligonucleotides, Oligosaccharides and Oligopeptides. Angew Chem Int Ed Engl 59, 19762–19772 (2020).

13. Gabay, M., Li, Y. & Felsher, D.W. MYC activation is a hallmark of cancer initiation and maintenance. Cold Spring Harb Perspect Med 4 (2014).

14. Dhanasekaran, R. et al. The MYC oncogene - the grand orchestrator of cancer growth and immune evasion. Nat Rev Clin Oncol 19, 23–36 (2022).

15. Han, H. et al. Small-Molecule MYC Inhibitors Suppress Tumor Growth and Enhance Immunotherapy. Cancer Cell 36, 483-497.e415 (2019).

16. Hammoudeh, D.I., Follis, A.V., Prochownik, E.V. & Metallo, S.J. Multiple independent binding sites for small-molecule inhibitors on the oncoprotein c-Myc. J Am Chem Soc 131, 7390–7401 (2009).

17. Jung, K.-Y. et al. Perturbation of the c-Myc–Max Protein–Protein Interaction via Synthetic α-Helix Mimetics. Journal of Medicinal Chemistry 58, 3002–3024 (2015).

18. Draeger, L.J. & Mullen, G.P. Interaction of the bHLH-zip domain of c-Myc with H1-type peptides. Characterization of helicity in the H1 peptides by NMR. J Biol Chem 269, 1785–1793 (1994).

19. Truica, M.I., Burns, M.C., Han, H. & Abdulkadir, S.A. Turning Up the Heat on MYC: Progress in Small-Molecule Inhibitors. Cancer Res 81, 248–253 (2021).

20. Bonger, K.M., Chen, L.C., Liu, C.W. & Wandless, T.J. Small-molecule displacement of a cryptic degron causes conditional protein degradation. Nat Chem Biol 7, 531–537 (2011).

21. Bonger, K.M., Rakhit, R., Payumo, A.Y., Chen, J.K. & Wandless, T.J. General method for regulating protein stability with light. ACS Chem Biol 9, 111–115 (2014).

22. Jin, J.W. et al. Development of an α-synuclein knockdown peptide and evaluation of its efficacy in Parkinson’s disease models. Communications Biology 4, 232 (2021).

23. Qu, J. et al. Specific Knockdown of α-Synuclein by Peptide-Directed Proteasome Degradation Rescued Its Associated Neurotoxicity. Cell Chem Biol 27, 751-762.e754 (2020).

24. Dang, C.V. MYC on the path to cancer. Cell 149, 22–35 (2012).

25. Madden, S.K., de Araujo, A.D., Gerhardt, M., Fairlie, D.P. & Mason, J.M. Taking the Myc out of cancer: toward therapeutic strategies to directly inhibit c-Myc. Molecular Cancer 20, 3 (2021).

26. Llombart, V. & Mansour, M.R. Therapeutic targeting of “undruggable” MYC. EBioMedicine 75, 103756 (2022).

27. Whitfield, J.R., Beaulieu, M.E. & Soucek, L. Strategies to Inhibit Myc and Their Clinical Applicability. Front Cell Dev Biol 5, 10 (2017).

28. Fletcher, S. & Prochownik, E.V. Small-molecule inhibitors of the Myc oncoprotein. Biochim Biophys Acta 1849, 525–543 (2015).

29. Berg, T. et al. Small-molecule antagonists of Myc/Max dimerization inhibit Myc-induced transformation of chicken embryo fibroblasts. Proc Natl Acad Sci U S A 99, 3830–3835 (2002).

30. Giorello, L. et al. Inhibition of cancer cell growth and c-Myc transcriptional activity by a c-Myc helix 1-type peptide fused to an internalization sequence. Cancer Res 58, 3654–3659 (1998).

31. Xie, D., Wang, F., Xiang, Y. & Huang, Y. Enhanced nuclear delivery of H1-S6A, F8A peptide by NrTP6-modified polymeric platform. Int J Pharm 580, 119224 (2020).

32. Ting, T.A., Chaumet, A. & Bard, F.A. Targeting c-Myc with a novel Peptide Nuclear Delivery Device. Scientific Reports 10, 17762 (2020).

33. Pescarolo, M.P. et al. A retro-inverso peptide homologous to helix 1 of c-Myc is a potent and specific inhibitor of proliferation in different cellular systems. Faseb j 15, 31–33 (2001).

34. Bidwell, G.L., 3rd & Raucher, D. Application of thermally responsive polypeptides directed against c-Myc transcriptional function for cancer therapy. Mol Cancer Ther 4, 1076–1085 (2005).

35. Bonger, K.M., Chen, L.-c., Liu, C.W. & Wandless, T.J. Small-molecule displacement of a cryptic degron causes conditional protein degradation. Nature Chemical Biology 7, 531–537 (2011).

36. Houston, K.M. et al. Development of β-Hairpin Peptides for the Measurement of SCF-Family E3 Ligase Activity in Vitro via Ornithine Ubiquitination. ACS Omega 2, 1198–1206 (2017).

37. Jin, J.W. et al. Development of an α-synuclein knockdown peptide and evaluation of its efficacy in Parkinson’s disease models. Commun Biol 4, 232 (2021).

38. Kammeyer, J.K., Blum, A.P., Adamiak, L., Hahn, M.E. & Gianneschi, N.C. Polymerization of Protecting-Group-Free Peptides via ROMP. Polym Chem 41, 3929–3933 (2013).

39. Berger, O. et al. Mussel Adhesive-Inspired Proteomimetic Polymer. Journal of the American Chemical Society 144, 4383–4392 (2022).

40. Bielawski, C.W. & Grubbs, R.H. Highly Efficient Ring-Opening Metathesis Polymerization (ROMP) Using New Ruthenium Catalysts Containing N-Heterocyclic Carbene Ligands C.B. is grateful to the National Science Foundation for a pre-doctoral fellowship. The authors thank Dr. Matthias Scholl for providing catalysts 4 a and 4c. Angew Chem Int Ed Engl 39, 2903–2906 (2000).

41. Bielawski, C.W. & Grubbs, R.H. Living ring-opening metathesis polymerization. Progress in Polymer Science 32, 1–29 (2007).

42. Callmann, C.E., Thompson, M.P. & Gianneschi, N.C. Poly(peptide): Synthesis, Structure, and Function of Peptide– Polymer Amphiphiles and Protein-like Polymers. Accounts of Chemical Research 53, 400–413 (2020).

43. Koren, E. & Torchilin, V.P. Cell-penetrating peptides: breaking through to the other side. Trends Mol Med 18, 385–393 (2012).

44. Behzadi, S. et al. Cellular uptake of nanoparticles: journey inside the cell. Chem Soc Rev 46, 4218–4244 (2017).

45. Hopewell, R. & Ziff, E.B. The nerve growth factor-responsive PC12 cell line does not express the Myc dimerization partner Max. Mol Cell Biol 15, 3470–3478 (1995).

46. Nalawansha, D.A., Li, K., Hines, J. & Crews, C.M. Hijacking Methyl Reader Proteins for Nuclear-Specific Protein Degradation. Journal of the American Chemical Society 144, 5594–5605 (2022).

47. Burslem, G.M. et al. The Advantages of Targeted Protein Degradation Over Inhibition: An RTK Case Study. Cell Chem Biol 25, 67-77.e63 (2018).

48. Bondeson, D.P. et al. Catalytic in vivo protein knockdown by small-molecule PROTACs. Nature Chemical Biology 11, 611–617 (2015).

49. Farrell, A.S. & Sears, R.C. MYC degradation. Cold Spring Harb Perspect Med 4 (2014).

50. Sears, R. et al. Multiple Ras-dependent phosphorylation pathways regulate Myc protein stability. Genes Dev 14, 2501–2514 (2000).

51. Zhou, Z., He, C. & Wang, J. Regulation mechanism of Fbxw7-related signaling pathways (Review). Oncol Rep 34, 2215–2224 (2015).

52. Zhang, X., Crowley, V.M., Wucherpfennig, T.G., Dix, M.M. & Cravatt, B.F. Electrophilic PROTACs that degrade nuclear proteins by engaging DCAF16. Nature Chemical Biology 15, 737–746 (2019).

53. Schneekloth, J.S., Jr. et al. Chemical genetic control of protein levels: selective in vivo targeted degradation. J Am Chem Soc 126, 3748–3754 (2004).

54. Lu, M. et al. Discovery of a Keap1-dependent peptide PROTAC to knockdown Tau by ubiquitination-proteasome degradation pathway. Eur J Med Chem 146, 251–259 (2018).

55. Ito, T., Yamaguchi, Y. & Handa, H. Exploiting ubiquitin ligase cereblon as a target for small-molecule compounds in medicine and chemical biology. Cell Chem Biol 28, 987–999 (2021).

56. Alabi, S.B. & Crews, C.M. Major advances in targeted protein degradation: PROTACs, LYTACs, and MADTACs. J Biol Chem 296, 100647 (2021).

57. Clark, G.J. et al. Peptides containing a consensus Ras binding sequence from Raf-1 and theGTPase activating protein NF1 inhibit Ras function. Proc Natl Acad Sci U S A 93, 1577–1581 (1996).

58. Pender, A. et al. Efficient Genotyping of KRAS Mutant Non-Small Cell Lung Cancer Using a Multiplexed Droplet Digital PCR Approach. PLoS One 10, e0139074 (2015).

59. Janssen, K. et al. Exploiting the intrinsic misfolding propensity of the KRAS oncoprotein. Proc Natl Acad Sci U S A 120, e2214921120 (2023).

60. Leinonen, H.M., Kansanen, E., Pölönen, P., Heinäniemi, M. & Levonen, A.L. Dysregulation of the Keap1-Nrf2 pathway in cancer. Biochem Soc Trans 43, 645–649 (2015).

61. Jin, J. et al. The peptide PROTAC modality: a novel strategy for targeted protein ubiquitination. Theranostics 10, 10141–10153 (2020).

62. Lou, K. et al. IFITM proteins assist cellular uptake of diverse linked chemotypes. Science 378, 1097–1104 (2022).

63. Saido, T. & Leissring, M.A. Proteolytic degradation of amyloid β-protein. Cold Spring Harb Perspect Med 2, a006379 (2012).

64. Su, L.C., Xu, W.D. & Huang, A.F. IRAK family in inflammatory autoimmune diseases. Autoimmun Rev 19, 102461 (2020).

65. Jiang, S.-Y. et al. Discovery of a potent HMG-CoA reductase degrader that eliminates statin-induced reductase accumulation and lowers cholesterol. Nature Communications 9, 5138 (2018).

