## Supplemental Information for "Heterobifunctional proteomimetic polymers for targeted protein degradation"

### TABLE OF CONTENTS

**List of Figures** ……………………………………………………………………………….….….…..... 3

**Materials and Methods** ……………………………………………………………………..….………. 4

I. Materials ……………………………………...……….……….….…………….…..... 4
II. Instrumentation ……….……………………………….………………...…….…..……… 4
III. Supplemental Methods ……...……………………….………………….……………...…… 5
IV. Supplemental Data …………………………………………………………………….. 11

**Supplemental Tables** ……….…………….…....…………………….……….………………….…… 20

**References** …………………………….……….………………….……………………….…...…… 22

### LIST OF FIGURES

**Fig. S1** Synthesis and characterization of H1 and RRRG conjugated norbornene monomers

**Fig. S2** $^1$H NMR spectra and polymerization kinetics

**Fig. S3** Determination of polymer molecular weights by SEC-MALSs & SDS-PAGE

**Fig. S4** Resistance of HYDRACs to enzymatic degradation

**Fig. S5** Circular dichroism spectrum of H1 homopolymer and scrambled control

**Fig. S6** Characterization of biotin terminating agent and quantification of termination efficiency

**Fig. S7** Characterization of Cy5.5-label and quantification of incorporation

**Fig. S8** Gating strategy for uptake analysis and inhibitor quantification

**Fig. S9** Gene expression profiles of HYDRAC vs scrambled controls or vehicle

**Fig. S10** Relative MYC mRNA levels measured by qPCR

**Fig. S11** Western blot of MYC protein levels in HEK293T cells

**Fig. S12** Representative western blot of MYC protein levels in PC3 cells

**Fig. S13** TMT-based whole proteome quantification

**Fig. S14** Body weights of mice treated with HYDRAC or vehicle

**Fig. S15** Characterization of VHL and KEAP norbornene monomers

**Fig. S16** Characterization of thalidomide norbornene monomers

**Fig. S17** $^1$H NMR spectra of VHL and KEAP1 homopolymers

**Fig. S18** Determination of $HYDRAC_{VHL, KEAP, CRBN}$ molecular weights by SEC-MALS

**Fig. S19** Characterization of RAS-targeting norbornene monomers

**Fig. S20** Western blot of KRAS protein levels in PANC-1 cells

**Supplementary Table 1** Calculated and obtained monomer weight values

**Supplementary Table 2** Summary Characterization of polymer formulations used

**Supplementary Table 3** Batch molecular weights for polymers used in each experiment

### Materials and Methods

#### I. Materials

Materials and reagents were purchased from commercial sources and used as received. Amino acids were purchased from AAPPTEC, ChemPep or NovaBiochem. N-(hexanoic acid)-cis-5-norbornene-exo-dicarboximide and modified second generation Grubbs' ruthenium initiator, (IMesH2)(C5H5N)2(Cl)2Ru=CHPh, were prepared as previously reported[1]. DMF-d7 from Sigma Aldrich (No. 1.11656) was used to monitor polymerizations via NMR. Protein gels (12% Mini-PROTEAN, 10 well, 30 ul) were purchased from BioRad (No. 4561043).

#### II. Instrumentation

**$^1$H Nuclear Magnetic Resonance ($^1$H NMR):** $^1$H NMR spectra were recorded on a 400 MHz Bruker Advance III HD Nanobay system equipped with SampleXpress autosampler in DMF-d7 at room temperature. Chemical shifts reported relative to proton signal of deuterated solvents.

**Mass Spectrometry (MS):** Electrospray Ionization Mass Spectrometry (ESI-MS) of products were collected using a Bruker Amazon-SL spectrometer configured with an ESI source in both negative and positive ionization modes. Analysis was performed at the Integrated Molecular Structure Education and Research Center (IMSERC) at Northwestern University.

**Analytical High-Performance Liquid Chromatography (HPLC):** Analytical HPLC analysis of peptides was performed on a Jupiter 4m Proteo 90Å Phenomenex column (150 x 4.60 mm) using a Hitachi-Elite LaChrom L-2130 pump with UV-Vis detector (Hitachi-Elite LaChrom L2420). Solvent system consists of (A) 0.1% TFA in water and (B) 0.1% TFA in acetonitrile.

**Preparative HPLC:** An Armen Glider CPC preparatory HPLC was used to purify all peptides. The solvent system consists of (A) 0.1% TFA in water and (B) 0.1% TFA in acetonitrile.

**Organic Phase Size Exclusion Chromatography coupled with Multi-angle Light Scattering (SEC-MALS):** Organic phase measurements were performed on a Phenomenex Phenogel 5μ, 1K-75K, 300 x 7.80 mm column in series with a Phenomex 5μ, 10K-1000K, 300 x 7.80 mm column with 0.05 M of LiBr in dimethylformamide (DMF) eluent. A Hitachi UV-Vis Detector L-2420, a Wyatt Optilab T-rEX refractive index detector operating at 658 nm and a Wyatt DAWN® HELEOS® II light scattering detector operating at 659 nm were used as detectors. Absolute molecular weight and polydispersity was calculated using the Wyatt ASTRA software with a dn/dc value of 0.179.

**Aqueous Phase SEC-MALS:** Aqueous phase SEC-MALS measurements were performed on an Agilent 1200 HPLC system with a PSS Suprema column using HPLC grade Phosphate Buffered Saline (PBS) buffer as the mobile phase. Detection consisted of a Wyatt Optilab T-rEX refractive index detector operating at 658 nm and a Wyatt DAWN® HELEOS® II light scattering detector operating at 659 nm. Absolute molecular weight and polydispersity was calculated using the Wyatt ASTRA software with a dn/dc value of 0.185.

**Fluorescence Plate Reader:** To determine Cy5.5 concentration/incorporation efficiency, an EnSpire Multimode Plate Reader (PerkinElmer) was used to quantify fluorescence levels.

**Confocal Laser Scanning Microscopy:** Imaging was done on a LEICA SP5 II laser scanning confocal microscope with a 100x oil immersion objective at 1.5x optical zoom. Z-stack images were taken with slice thickness at 0.26 µm and scan size of 1024 x 1024 pixels at 400 Hz. The cell nuclei (stained with DAPI) were imaged using a 358 nm laser with a 15% laser power.

### III. Supplemental Methods

#### Synthesis of Peptide Monomer (Nor-Peptide)

Peptides were synthesized on a Liberty Blue (CEM) automated microwave peptide synthesizer by solid phase peptide synthesis (SPPS) using standard FMOC chemistry. Peptides were subsequently capped with a norbornene–amino hexanoic acid, as previously described[1-3]. Briefly, amide coupling to N-(hexanoic acid)-cis-5-norbornene-exo-dicarboximide (3.0 equiv) was done in the presence of HBTU (2.9 equiv) and DIPEA (6.0 equiv) on resin. Cleavage from the resin was performed using 92.5:2.5:2.5:2.5 (% v/v) trifluoroacetic acid (TFA), triisopropyl silane, water, and DTT stirring at RT for 2 h. Following filtration and rotary evaporation of TFA, cleaved peptides were precipitated in cold diethyl ether, centrifuged down into a pellet, and dried under vacuum. This was then purified with a Jupiter Proteo 90 Å Phenomenex column (2050 × 25.0 mm) on an Armen Glider CPC preparatory-phase HPLC using a buffer B (acetonitrile with 0.1% TFA) gradient over 30 min to yield 90–95% purity, confirmed by analytical HPLC. Pure products were then analyzed by electrospray ionization mass spectroscopy on a Bruker amazon SL to confirm molecular weight (Supplementary Fig. 1). See Supplementary Table 1 for summary of peptide monomer sequences.

Nor-“Peptide”

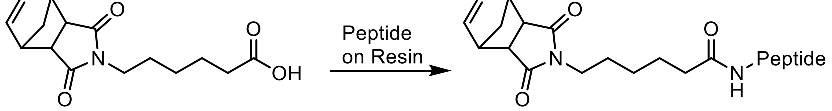


#### Synthesis of Cy5.5 Monomer

Norbornene-Cy5.5 (Nor-Cy5.5) was synthesized as described previously[4]. Briefly, 0179 mmol (1 equiv) of Cy5.5 was activated with 0.0179 mmol (1 equiv) of HBTU and 0.1076 mmol of DIPEA (6 equiv.) in DMF for 10 min. To this, 0.0538 mmol (3 equiv.) of 2-(2-aminoethyl)-3a,4,7,7a-tetrahydro-1H-4,7-methanoisoindole-1,3(2H)-dione was added followed by room temperature stirring for 2 h. The product was precipitated in cold diethyl ether, dissolved in a mixture of water and acetonitrile in 0.1% TFA and purified by reverse phase HPLC using a gradient 10%-60% acetonitrile in 0.1% TFA. Pure fractions were lyophilized. See Supplementary Fig. 7.

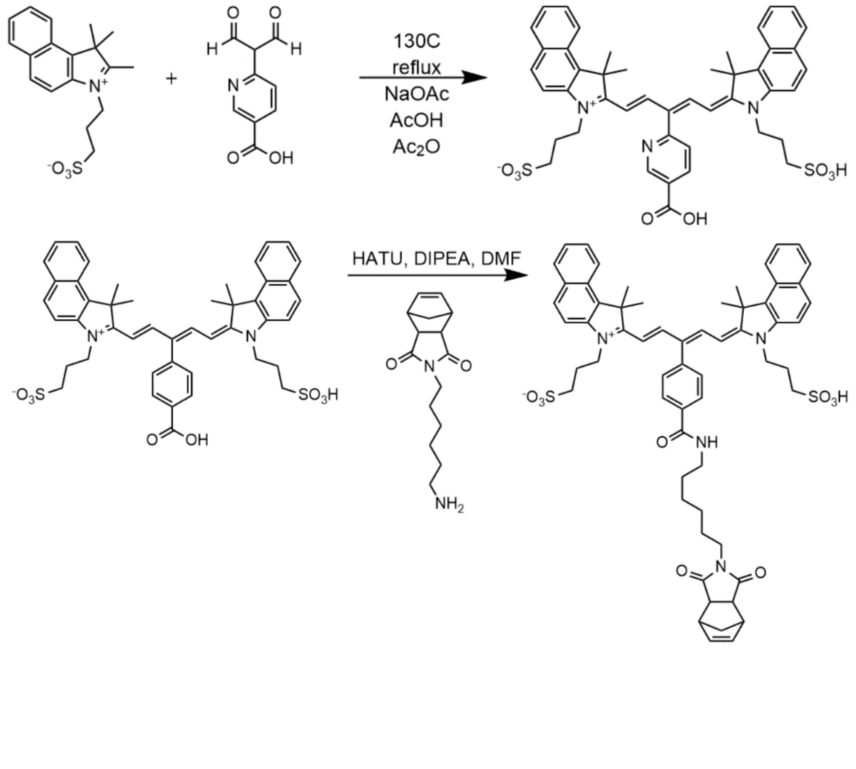

**Synthesis of Biotin Terminating Agent**

A biotin terminating agent was synthesized as previously reported[5]. Briefly, to a solution of NHS-biotin (215 mg, 0.88 mmol) in dry dimethylformamide (3 ml), HBTU (215 mg, 0.88 mmol), 1-Hydroxybenzotriazole (HOBt, 118.8 mg, 0.88 mmol), and triethanolamine (TEA, 122.7 μL, 0.88 mmol) were added. The diamine (124 mg, 0.38 mmol) was added to the solution in dimethylformamide (3 ml) and stirred under inert gas for two days and precipitated into cold ether and centrifuged. The precipitate was collected via filtration and recrystallized in toluene and methanol (4:1). The resulting product was filtered and collected as a white powder and characterized by ESI-MS. ESI-MS expected $[M+H]^+$: 778.35, found 779.47. $^1H$ NMR (500Hz DMSO) δ 1.20-1.67 (m, 12H), 2.03 (t, *J* = 7.3 Hz, 2H), 2.2 (t, *J* = 7.4 Hz, 2H), 2.53-2.67 (m, 6H), 2.78-2.85 (m, 2H), 3.04-3.13 (m, 2H), 3.18-3.25 (m, 4H), 4.09-4.16 (m, 2H), 4.26-4.34 (m, 2H), 4.64-4.71 (d, *J* = 3.8 Hz, 4H), 5.84 (t, *J* = 3.8 Hz, 2H), 6.35 (s, 2H), 6.41 (s, 2H), 6.87 (d, *J* = 8.6 Hz, 4H), 7.10 (d, *J* = 8.7 Hz, 4H), 7.81 (t, *J* = 5.6 Hz, 2H). See Supplementary Fig. 5.

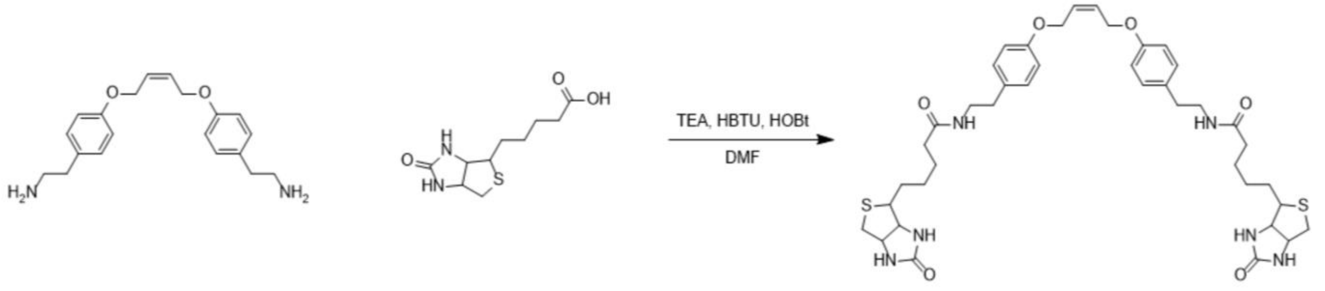


**Synthesis of Thalidomide Monomer**

494.4 mg (2.1 equiv) t-butyl bromoacetate, 200 mg (1 equiv) potassium iodide, 910.8 mg (9 equiv) sodium bicarbonate, and 495.8 g 4-hydroxy thalidomide (1.5 equiv) were mixed in dry DMF. The reaction vessel was lowered into a 65 °C oil bath and nitrogen flow was set up. The reaction was run overnight, filtered, and rotary evaporated. The remaining reaction mixture was precipitated into cold diethyl ether, centrifuged down into a pellet, and dried under vacuum. To remove Boc-protecting group, 10 mL of 90% TFA in DCM was added for 4 h. This was precipitated into cold diethyl ether, centrifuged down into a pellet, and dried under vacuum. Lastly, 400 mg (1 equiv) Boc-deprotected intermediate product, 2.5 mL DIPEA (12 equiv), 272.8 mg EDC (1.2 equiv), and 163.8 mg (1.2 equiv) N-hydroxyl succinimide were added to dry DMF at room temperature and stirred for 3 h. Next, 622.3 mg (2 equiv) of norbornene-amine with a six-carbon spacer was added and stirred overnight. The mixture was rotary evaporated and aziotroped with toluene 5 times. The final product was purified using preparatory HPLC and pure fractions were lyophilized. ESI-MS expected $[M+H]^+$: 576.22, found 577.30. $^1H$ NMR (600Hz, DMSO) δ 0.95-1.49 (m, 10H, $CH_2$), 1.98-2.08 (m, 1H, NC-CH), 2.51-2.64 (m, 2H, O=C-CH, NC-CH), 2.64-2.71 (br s, 2H, N-CH), 2.83-2.95 (m, 2H, O=C-CH), 3.04-3.17 (m, 4H, CH), 3.27-3.40 (m, 2H, N-CH), 4.71-4.80 (s, 2H, O-CH), 5.06-5.16 (m, 1H, N-CH), 6.22-6.36 (m, 2H, C=CH), 7.34-7.42 (d, *J* = 8.4 Hz, 1H, Ar), 7.45-7.52 (d, *J* = 7.2 Hz, 1H, Ar), 7.77-7.84 (t, *J* = 5.3 Hz, 1H, Ar), 7.88-7.95 (m, 1H, CO-NH), 11.01-11.24, (s, 1H, CO-NH). $^{13}C$ NMR (151 MHz, DMSO) δ 21.99, 25.74, 25.99, 27.13, 28.78, 30.95, 37.80, 38.15, 39.10, 39.24, 39.38, 39.52, 39.66, 39.80, 39.94, 42.33, 44.45, 47.22, 48.81, 67.65, 116.04, 116.83, 120.38, 133.03, 136.93, 137.63, 155.05, 165.51, 166.63, 166.74, 169.87, 172.78, 177.69. See Supplementary Fig. 15.

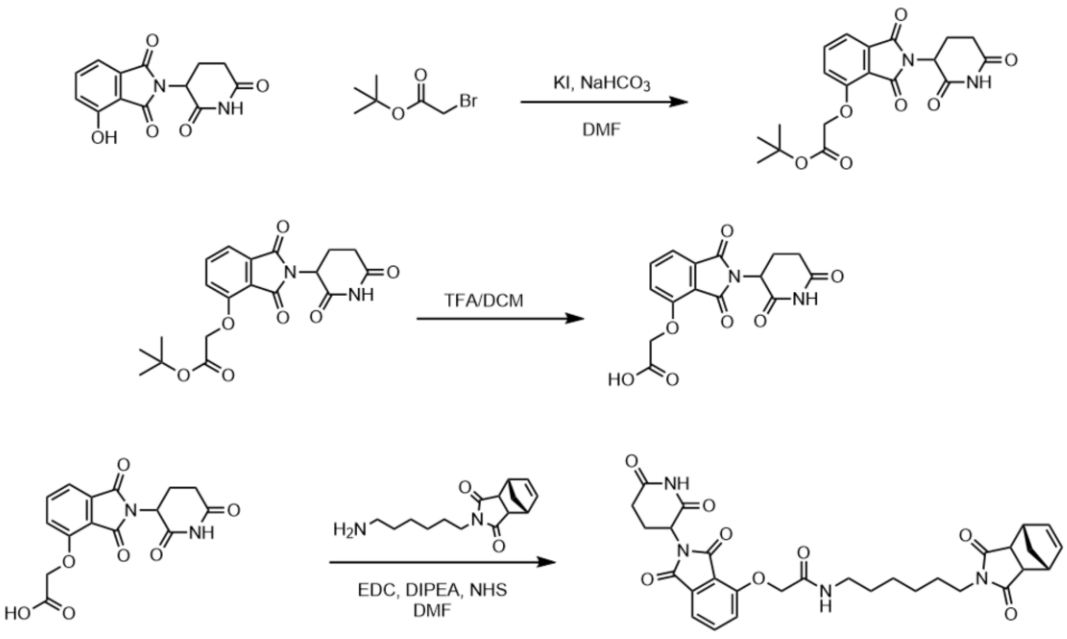


**Polymerization**

Polymerization reactions were carried out in a glove box under flowing nitrogen gas, as previously described [1-3]. $^1$H NMR spectroscopy was used to track polymerization kinetics and confirm monomer consumption (Supplementary Fig. 2). Copolymer formulations with both H1 and RRRG side chains were made by adding equivalent molarities of the two norbornene monomers at the onset of the polymerization due to their similar reaction kinetics. A library of compounds was synthesized with differing H1:RRRG or H1:E3 ligase ratios (Supplementary Table 2). Polymers were terminated with ethyl vinyl ether (10 equiv) for 1 h at room temperature or biotin termination agent (5 equiv) for 24h at room temperature, then precipitated, washed with cold diethyl ether 3 times, and collected by centrifugation. Molecular weight and polydispersity were assessed by SEC-MALS at 65°C in 0.05 M LiBr in DMF or SDS-PAGE gel (Supplementary Fig. 3).

**Evaluation of Proteolytic Stability**

In a typical cleavage experiment, 0.1 μM Chymotrypsin was incubated with different concentrations of polymers dissolved in 2 mL of PBS. Into this, 2 μL of a chymotrypsin stock solution was added and stirred in a preheated oil bath at 37° C. Aliquots of the reaction were taken for HPLC quantification of proteolysis at predetermined time points and the experiment repeated a minimum of three times.

**Circular Dichroism Spectroscopy**

CD spectra were obtained on a JASCO J-815 spectrometer equipped with a Peltier sample holder for temperature control. Polymer samples were dissolved in buffered solvent to concentrations of approximately 1 μM, and placed into a 1 cm quartz cuvette (Hellma). Protein samples were measured at 5 μM. All samples were equilibrated at room temperature for at least thirty minutes prior to measurement, with mixtures (i.e. protein and polymer combinations) being generally equilibrated for 12 hours prior to measurement. Spectra were obtained using a scan speed of 10 nm/min, using a 1 nm bandwidth, 4 sec response time, and with three accumulations. Background spectra of solvent were obtained prior to measurement of the sample solutions and subtracted from the sample spectra using JASCO Spectra Analysis. Spectra were cut off where the HT voltage exceeded 750 V. For melting experiments, the temperature was varied between 20 and 91 ºC with steps of 1 ºC/min, while measuring the ellipticity at 222 nm with a 2 nm bandwidth.

**Cell Culture**

PC-3, PC-12, A549 cells were obtained from ATCC; Mouse MycCaP cells were the kind gift of Charles Sawyers (Memorial Sloan-Kettering Cancer Centre). All cells were authenticated and tested as mycoplasma-free. PC-3 and A549 cells were grown in F12K (Gibco) medium; MycCaP cells in DMEM medium (Gibco), all supplemented with 10% heat-inactivated fetal bovine serum (FBS, Gibco). PC-12 cells were grown in F-12K medium (ATCC) with 5% heat-inactivated FBS and 10% heat-inactivated horse serum (Thermo). All cells were cultured in 1% Penicillin-Streptomycin (10,000U/ml, Life Technologies) and 5% CO2 in a humidified incubator at 37 °C.

**Confocal Laser Scanning Microscopy (CLSM):** Imaging was done on a LEICA SP5 II laser scanning confocal microscope with a 100x oil immersion objective at 1.5x optical zoom. Z-stack images were taken with slice thickness at 0.26 µm and scan size of 1024 x 1024 pixels at 400 Hz. The cell nuclei (stained with Hoechst 3342) were imaged using a 358 nm laser with a 15% laser power. Cell membrane imaging was done using WGA-488.

**Cell Proliferation Assays**

Cell viability was estimated using the CellTiter-Glo® Luminescent Cell Viability Assay (Promega). According to cell type and experimental setting, 1000 to 5000 cells/well were seeded in 96 well white plates with clear bottom (ThermoFisher, 136101) and allowed to adhere overnight. After 3 to 5 days following the treatment, CellTiter-Glo reagent was added mixed 1:1 vol/vol with fresh media, plates incubated for 10 minutes on a shaking rotator followed by 10 minutes without shaking at room temperature (protected from light using aluminum foil) and luminescence was measured using plate reader (Perkin Elmer Victor 3V).

**Western Blot analysis**

Cells were lysed in RIPA (Sigma) lysis buffer containing 1x Halt™ Protease and Phosphatase Inhibitor Cocktail (ThermoFisher). Protein concentration was measured by Bio-Rad Bradford reagent. Protein samples were prepared by addition of 4x Laemmli Sample buffer (Bio-Rad) and 2-mercaptoethanol (Bio-Rad) and resolved on 4-12% SDS-PAGE (Sodium dodecyl sulfate–polyacrylamide) gels, which were subsequently transferred to PVDF membranes (Bio-Rad) using Trans-Blot Turbo transfer buffer (Bio-Rad) and a Trans-Blot Turbo Transfer System (Bio-Rad). Membranes were blocked for 1 h at room temperature with 5% blotting-grade blocker non-fat dry milk (Bio-Rad), followed by overnight 4 °C incubation with the appropriate primary antibody and 1 h room temperature incubation with an anti-rabbit or anti-mouse IgG (H + L)-HRP conjugate (Bio-Rad) secondary antibody. Blots were imaged using Supersignal West Femto Maximum Sensitivity Substrate detection system (Thermo) and the ChemiDoc Imaging System (Bio-Rad). The following primary antibodies were used: c-Myc (Y69) (Abcam #ab32072), Streptavidin-HRP (PerkinElmer NEL750), GAPDH (Cell Signaling #3683) and Actin (Cell Signaling #5125). Quantification analyses were performed by Biorad ChemiDoc Imager and Image Lab software.

**In vitro pull down assay**

To confirm HYDRAC direct binding to endogenous MYC protein in cell lysate complex, biotin conjugated polymers were synthesized. Collected PC3 cell pellet (2 million cells per pulldown condition) was suspended in 300 µL pulldown lysis buffer (pH 7.4 50mM Tris-HCl, 150 mM NaCl,

1mM EDTA, 0.1% IGEPAL nonionic detergent + 1x Halt protease/phosphatase inhibitor cocktail) and lysed by repeat (3x) freeze-thawing in liquid nitrogen. The lysed cell pellets were spun at 4 °C for 15 minutes at 15,000 rpm on a Eppendorf 5424R centrifuge. The collected supernatants were pre-cleared with Pierce streptavidin magnetic beads (Thermo, 88817) for 1 h at 4 °C. Around 300 µg was applied to each sample and incubated with 5 µM or 10 µM of various polymer compositions or D-Biotin on a rotator over night at 4 °C. Next day, 60 µl of streptavidin beads were added to each sample and further rotated for 1 h at 4 °C. Beads were washed with wash buffer (lysis buffer containing 0.1% BSA) for 3 times and another 3 times with lysis buffer, then eluted with 50 µL of 2x Laemmli sample buffer and boiled at 95 °C for 5 min. The supernatant was subjected to Western Blot.

**RNA-sequencing**

PC-3 cells were treated with 5µM HYDRAC or 5 µM scramble control for 24 hours. Cells were collected using trypsinization and washed with ice-cold PBS and later centrifuged at 1500 rpm for 5 minutes. Total RNA was extracted from cell pellet using Qiagen RNeasy Plus kit. The stranded total RNA-seq was conducted by the Northwestern University NUSeq Core Facility. Briefly, total RNA examples were checked for quality on Agilent Bioanalyzer 2100 and quantity with Qubit fluorometer. The NEBNext Ultra II RNA Library Prep Kit for Illumina was used to prepare sequencing libraries. The Kit procedure was performed without modifications. This procedure includes rRNA depletion, remaining RNA purification and fragmentation, cDNA synthesis, end preparation, Illumina adapter ligation, library PCR amplification and validation. Illumina HiSeq 4000 Sequencer was used to sequence the libraries with the production of single-end, 50 bp reads.

**Mass spectrometry-based whole proteome analysis.**

PC3 cells were treated with indicated polymer formulations for 24 hours. After treatment, the cells were collected and lysed using a probe sonicator (40%, 3, 5 pulses) in 150 µL of lysis buffer (PBS supplemented with Roche complete protease inhibitor cocktail). The protein concentration in the lysate was determined using a DC assay and adjusted to 1.0 mg/mL. Next, 100 µL of protein samples containing 100 µg of protein were transferred to new Eppendorf tubes (1.5 mL), to which 48 mg of urea was added (final concentration is 8 M). Subsequently, 5 µL of DTT (200 mM stock in water) was added to the tubes to achieve a final concentration of 10 mM. The samples were then incubated at 65 °C for 15 min. Afterward, 5 µL of iodoacetamide (400 mM stock in water) was added to the tubes to achieve a final concentration of 20 mM. The samples were then incubated in the dark at 37 °C for 30 min. Next, 600 µL of MeOH, 200 µL of CHCl3, and 500 µL of water were added to the tubes, mixed, and then centrifuged at 10,000 g for 10 min at 4 °C. A protein disc was formed at the interface of the CHCl3 and aqueous layers. The top layer was aspirated, and 1 mL of MeOH was added. The samples were then pelleted after centrifugation (10,000 g, 10 min, 4 °C). The protein pellets were resuspended in 160 µL of EPPS buffer (200 mM, pH 8). Next, 4 µL of LysC solution (0.5 µg/µL in water) was added to each sample, and the samples were incubated at 37 °C for 2 h. Afterward, 10 µL of trypsin (0.5 µg/µL in trypsin buffer) and 1.8 µL of CaCl2 (100 mM stock in water) were added to each sample, and the samples were incubated at 37 °C for 12 h. The peptide concentration was determined using a micro BCA assay kit. For each sample, a volume corresponding to 12.5 µg of peptides was transferred to a new Eppendorf tube, and the total volume was adjusted to 35 µL using EPPS buffer (200 mM, pH 8). Next, 9 µL of CH3CN was added to each sample, followed by the addition of 6 µL of TMT tags (20 µg/µL in dry CH3CN). The samples were then incubated at room temperature for 1 hour. To quench the TMT labeling reaction, 6 µL of hydroxylamine (5% in water) was added to each sample. After incubating at room temperature for 15 minutes, 2.5 µL of formic acid was added. The samples were combined into a new Eppendorf tube and dried using a SpeedVac. Subsequently, the peptides were fractionated into 12 fractions using the Thermo Vanquish UHPLC fractionator. The peptides were analyzed using liquid chromatography tandem mass-spectrometry on an Orbitrap Eclipse Tribrid Mass Spectrometer coupled to a Vanquish Neo UHPLC System. Peptides were injected onto an EASY-Spray HPLC column (C18, 2 µm particle size, 75 µm inner diameter, 250 mm length) and separated at a flow rate of 0.25 µL/min using the following gradient: 5% buffer B (80% acetonitrile with 0.1% formic acid) in buffer A (water with 0.1% formic acid) from 0-15 min, 5-45% buffer B from 15-155 min, and 45-100% buffer B from 155-180 min. The nano-LC electrospray ionization source was set to a voltage of 1.5 kV. The scan sequence began with an MS1 master scan (Orbitrap analysis, resolution 120,000, 375-1600 m/z, RF lens 30%, standard AGC target, auto maximum injection time). The top ten precursors were then selected for MS2/MS3 analysis. MS2 analysis involved quadrupole isolation (isolation window 1.2) of precursor ion, followed by HCD collision entry in the ion trap (standard AGC, collision energy 32%, maximum injection time 35 ms). Following the acquisition of each MS2 spectrum, synchronous precursor selection (SPS) enabled the selection of up to 10 MS2 fragment ions for MS3 analysis. The MS3 precursors were fragmented by HCD and analyzed using the Orbitrap (collision energy 55%, AGC 250%, maximum injection time 200 ms, resolution was 50,000). The RAW data was searched in Proteome Discoverer 2.5.

**In vivo studies**

All animal experiments and procedures were performed in compliance with ethical standards and the approval of Northwestern University Animal Care and Use Committee (IACUC). FVB mice were obtained from Jackson Laboratory. All the mice used in this study were maintained in a pathogen-free animal barrier facility. All the experiments were initiated with mice of age 6 to 8 weeks. For allograft models, MYC-driven murine prostate cancer MycCaP cells were suspended at a concentration of 10 million cells/mL in 50%Matrigel-50%PBS and 100 µL of this solution for a total of 1 million cells subcutaneously injected into the flanks of mice. Mice were grouped and treatment was started when the tumor size reached around 150 to 180 $mm^3$. Polymers were dissolved in 10% DMSO (Sigma) in PBS for I.P. administration.

**Immunofluorescence and Immunohistochemistry**

Tissues were fixed in 10% neutral buffered formalin for 48 h at 4 °C and transferred to 70% ethanol before paraffin processing at the Northwestern University Mouse Histology and Phenotyping Laboratory (MHPL) histology core. Primary antibodies used are: c-Myc (Y69) (Abcam #ab32072), Ki-67 (Abcam #ab15580), Cleaved Caspase-3 (Cell Signaling #9661). For Immuno-histochemistry, slides were incubated with ImmPRESS HRP anti-mouse (Vector #MP-7402) or ImmPRESS HRP anti-rabbit (Vector#MP-7401). Expression was visualized by using AEC peroxidase substrate (Vector #SK-4200). Slides were incubated with Hematoxylin (Vector #3404) and mounted with Glycergel Mounting Medium (Dako #C056330-2). For immuno-fluorescence, slides were incubated with secondary antibodies labelled with Alexa Fluor 488 anti-rabbit (Thermo Scientific # A11008), Alexa 594 anti-rabbit (Thermo Scientific #A-21207), Alexa 594 anti-mouse (Molecular probes #A11005) and Alexa Fluor 568 Goat anti-Mouse IgG (H+L) (Thermo Scientific #A-11004). Slides were counterstained with DAPI (Sigma #D-9542) and mounted with ProLong Gold Antifade reagent (Invitrogen/Molecular Probes #P36961). Immunofluorescence images were visualized using fluorescent microscope or Leica A1R spectral confocal microscope.

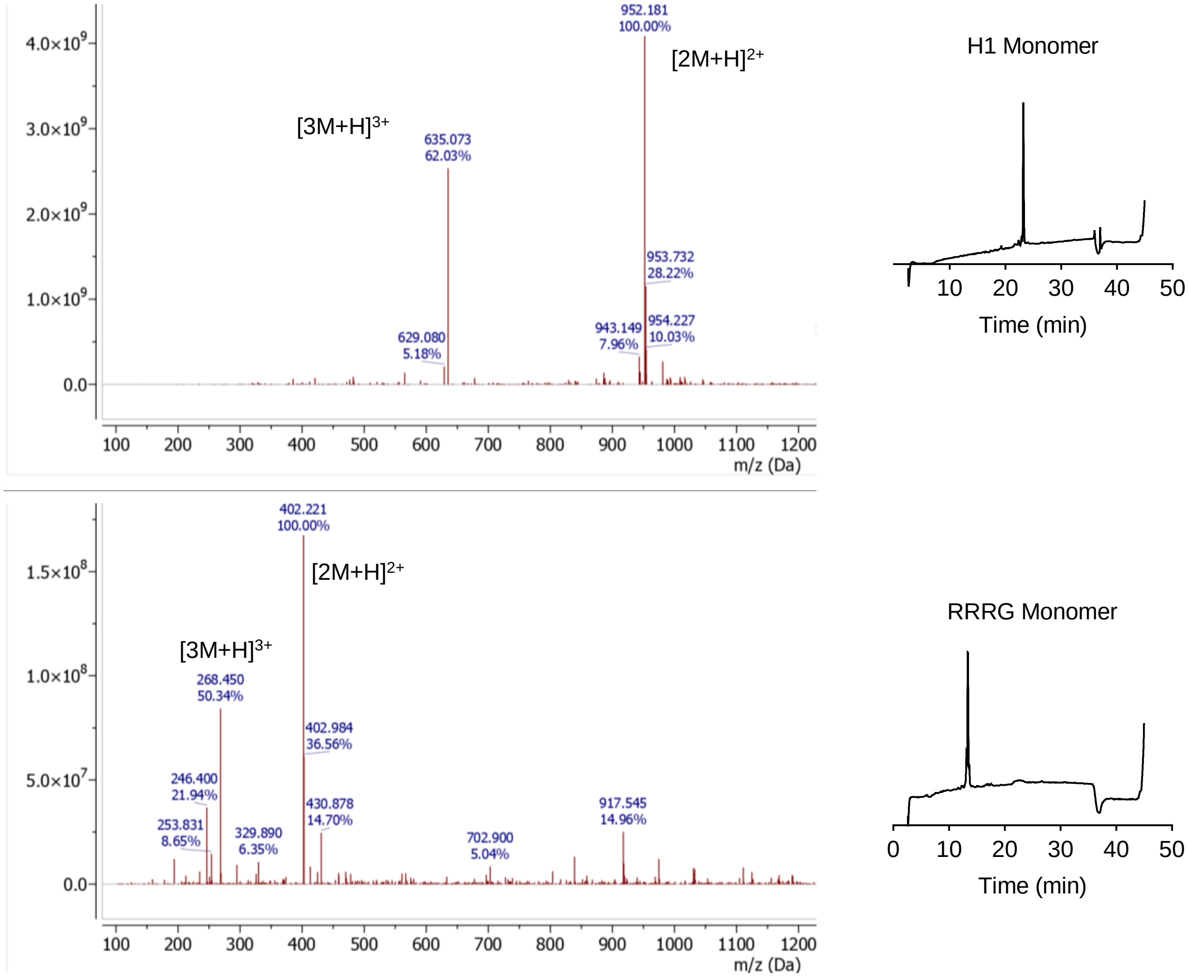


**Supplementary Fig. 1 | Synthesis and characterization of H1 (NELKRAFAALRDQI) and RRRG conjugated norbornene-based monomers.** ESI-MS and analytical HPLC of purified monomers at a 15-65% ACN over 30 min gradient. H1 monomer ESI-MS expected $[2M+H]^{2+}$: 952.1, found: 952.6. RRRG monomer ESI-MS expected $[2M+H]^{2+}$: 401.5, found: 401.6

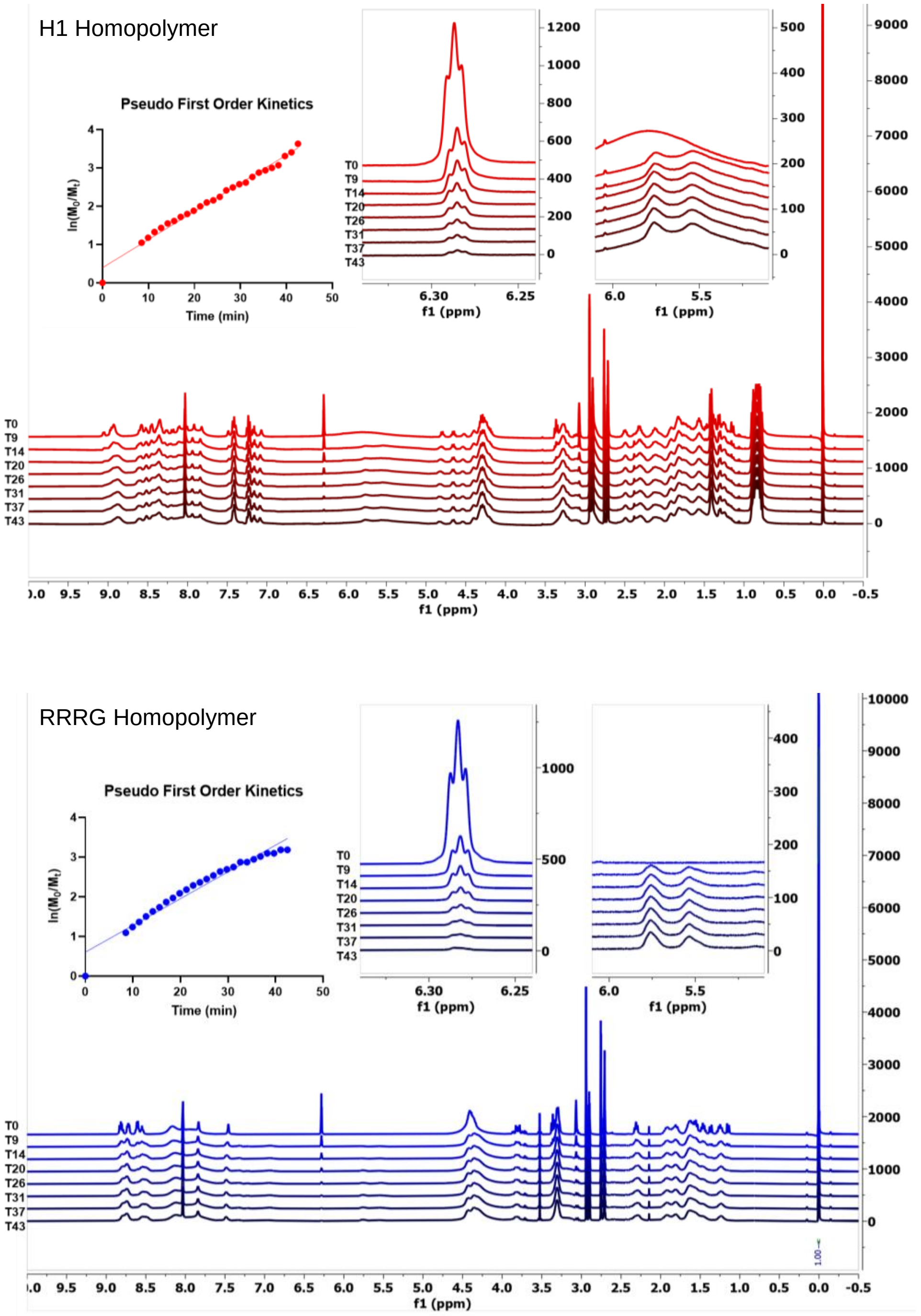


**Supplementary Fig. 2 | Representative $^{1}H$ NMR spectra and polymerization kinetic analysis.** Resonance at δ 6.3 ppm corresponds to monomer olefin protons. Final spectrum recorded at the end of polymerization shows consumption of the monomer. Resonances at ~ δ 5.6 pp m correspond to cis/trans olefinic protons of the polymerized material.

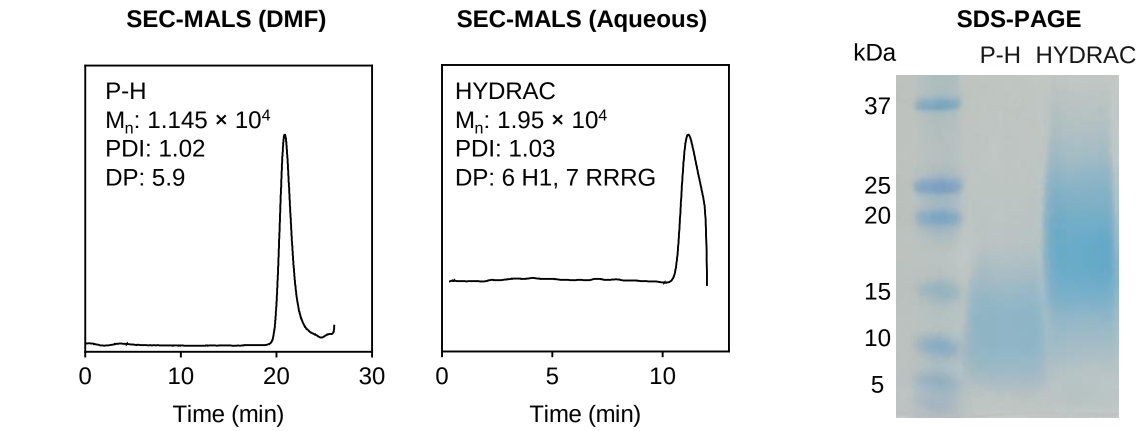


**Supplementary Fig. 3 | Determination of polymer molecular weights by SEC-MALS & SDS-PAGE.** Representative differential refractive index (dRI) chromatograms and SDS-PAGE gels of H1 homopolymer (P-H) and HYDRACs shown.

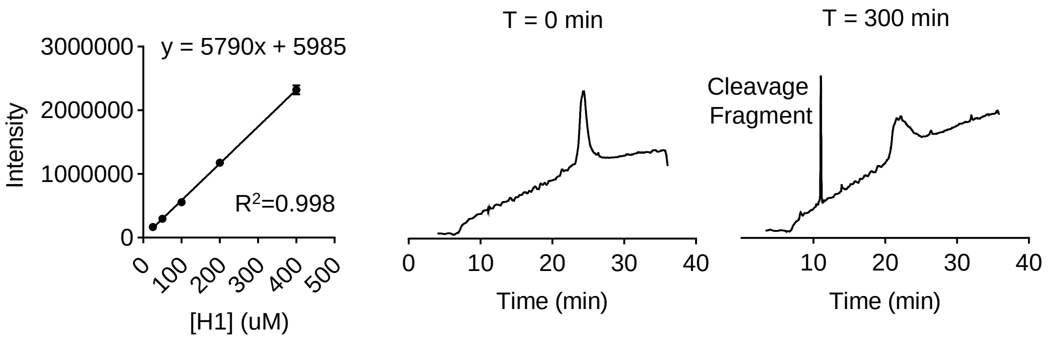


**Supplementary Fig. 4 | Resistance of HYDRACs to enzymatic degradation.** Representative HPLC traces of HYDRACs following incubation with 0.1 µM chymotrypsin over time (right) and H1 fragment calibration curve (left) used to quantify enzyme kinetics. Cleavage rates of H1 calculated using HPLC at a 15-65% ACN over 30 min gradient and shown in main figure 1d.

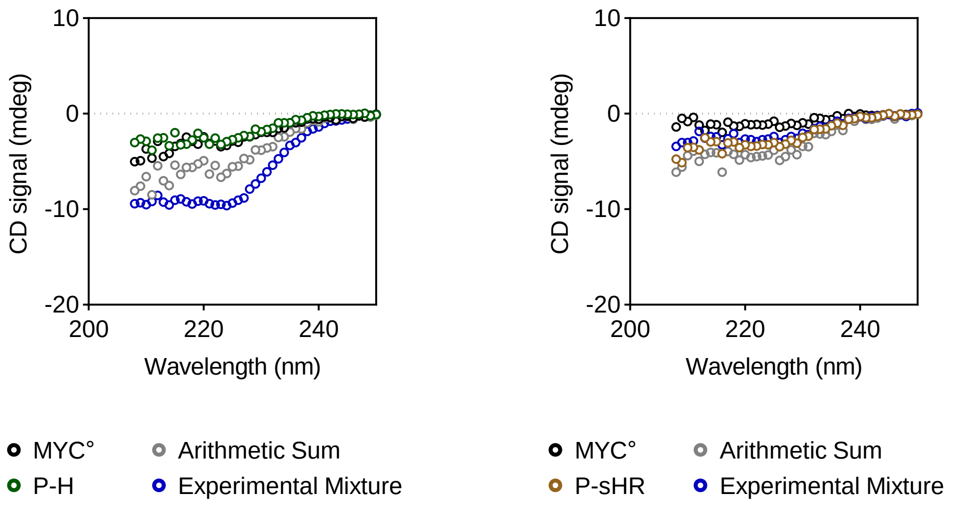


**Supplementary Fig. 5 | Circular Dichroism spectrum of H1 homopolymer and scrambled control.** Far-ultraviolet (UV) CD spectra of the b-HLH-LZ domain of MYC (MYC°) (black, 5 µM monomer units), P-H (green, 1 µM polymer units), P-sHR (brown, 1 µM polymer units), the arithmetic sum (gray), and the spectrum of a mixture of either polymer plus MYC° at a 1 to 5 molar ratio, equivalent to 1 to 1 binding peptide to protein ratio (blue) recorded at 20° C.

**a**

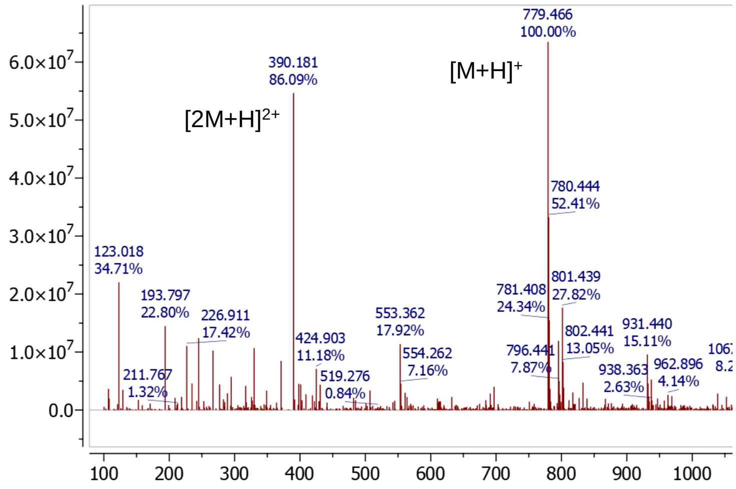


**b**

| | $M_n$ | Biotin Incorporation |
|---|---|---|
| P-$H_B$ | $1.1 \times 10^4$ | 98.8% |
| $HYDRAC_B$ | $1.7 \times 10^4$ | 97.3% |
| P-$sH_B$ | $1.0 \times 10^4$ | 92.4% |
| P-$sHR_B$ | $1.8 \times 10^4$ | 95.6% |

**Supplementary Fig. 6 | Characterization of biotin terminating agent and quantification of termination efficiency. a,** Characterization of biotin terminating agent via ESI-MS. ESI-MS expected $[M+H]^+$: 778.35, found 779.47. **b,** Determination of biotin incorporation efficiency as quantified by the QuantTag Kit based on manufacturer protocol (Vector Labs, BDK-2000).

**a**

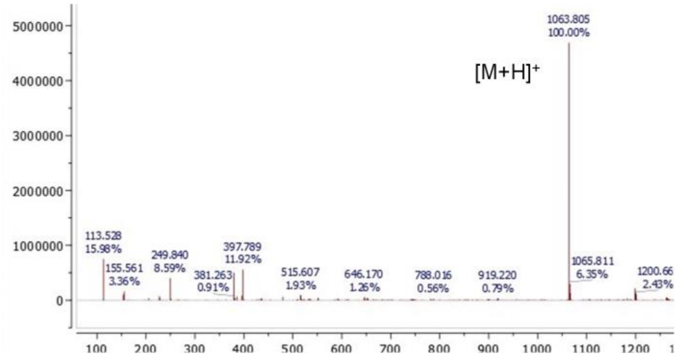


**b**

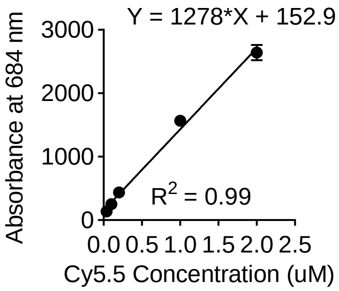


**c**

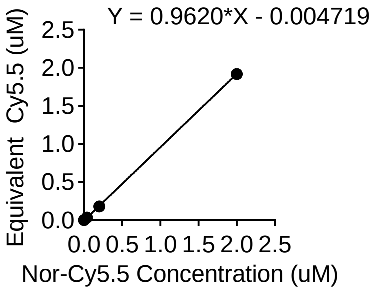


**d**

| | $M_n$ | Cy5.5 Incorporation |
|---|---|---|
| P-$H_F$ | $1.06 \times 10^4$ | 94.4% |
| P-$R_F$ | $8.60 \times 10^3$ | 87.2% |
| $HYDRAC_F$ | $1.70 \times 10^4$ | 96.2% |
| P-$sHR_F$ | $1.60 \times 10^4$ | 92.4% |

**Supplementary Fig. 7 | Characterization of Cy5.5-label and quantification of incorporation. a,** ESI-MS of Nor-Cy5.5 monomer added to terminus of polymer for fluorescent labeling. ESI-MS expected $[M+H]^+$: 1064.35, found 1063.8. **b,** Absorbance calibration curve of Cyanine5.5 carboxylic acid used to quantify Cy5.5 incorporation. Data recorded at 684 nm. **c,** Characterization of Nor-Cy5.5 absorbance as a function of Cy5.5 levels. **d,** Quantification of Cy5.5 incorporation for labeled polymers. $M_n$ of a reaction aliquot was determined by SEC-MALS prior to addition of one equivalent Nor-Cy5.5. Incorporation percentage determined by comparing expected absorbance levels to experimentally obtained values in n = 3 different technical replicates.

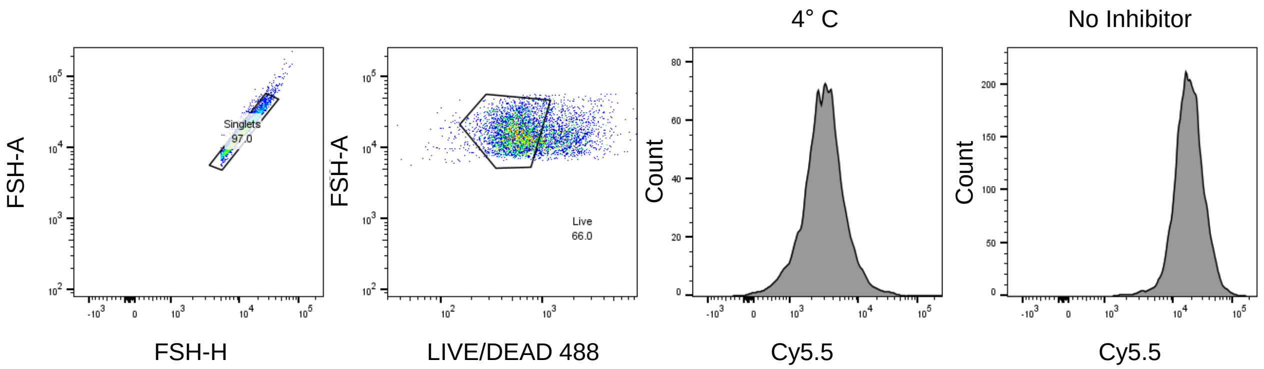


**Supplementary Fig. 8 | Gating strategy used for uptake analysis and inhibitor quantification.** Live cells were selected from the singlet population and Cy5.5 signal assessed.

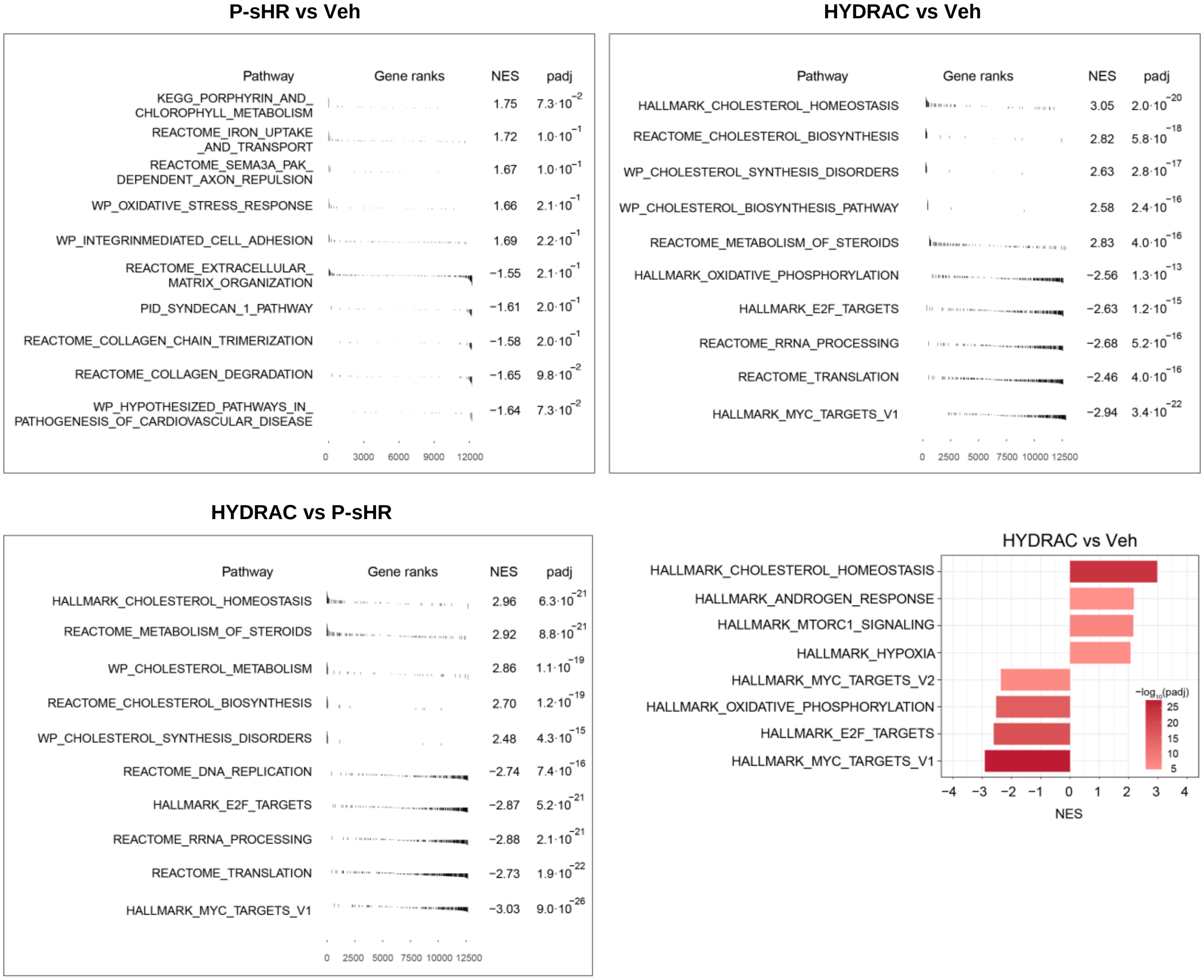


**Supplementary Fig. 9 | Gene expression profiles of HYDRAC vs P-sHR or vehicle.** List of gene expression profiles of scramble control or vehicle versus HYDRAC-treated PC3 cells. Normalized enrichment scores (NES) and p values of top gene sets are listed.

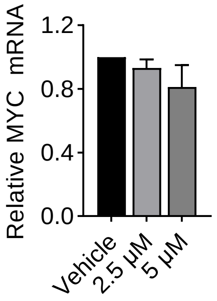


**Supplementary Fig. 10 | HYDRAC treatment has no noticeable effects on MYC mRNA levels.** PC3 cells treated with 2.5 or 5 µM HYDRACs with mRNA levels quantified via qPCR.

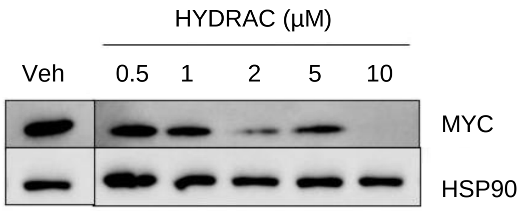


**Supplementary Fig. 11 | Western blot of MYC protein levels in HEK293T Cells.** HEK293T cells were treated with HYDRACs at increasing concentrations for 24 h and MYC protein levels tracked by Western Blot.

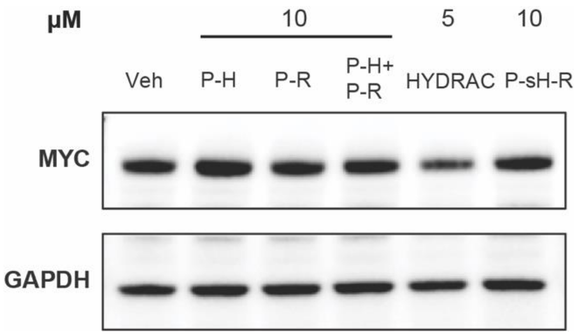


**Supplementary Fig. 12 | Representative western blot of MYC protein levels in PC3 cells.** Endogenous MYC levels in PC3 cells following 24 h incubation with indicated polymer formulations at 10 µM (P-H, P-R, P-H + P-R, P-sHR) or 5 µM HYDRAC. P-H + P-R indicates independent addition of both polymers to media.

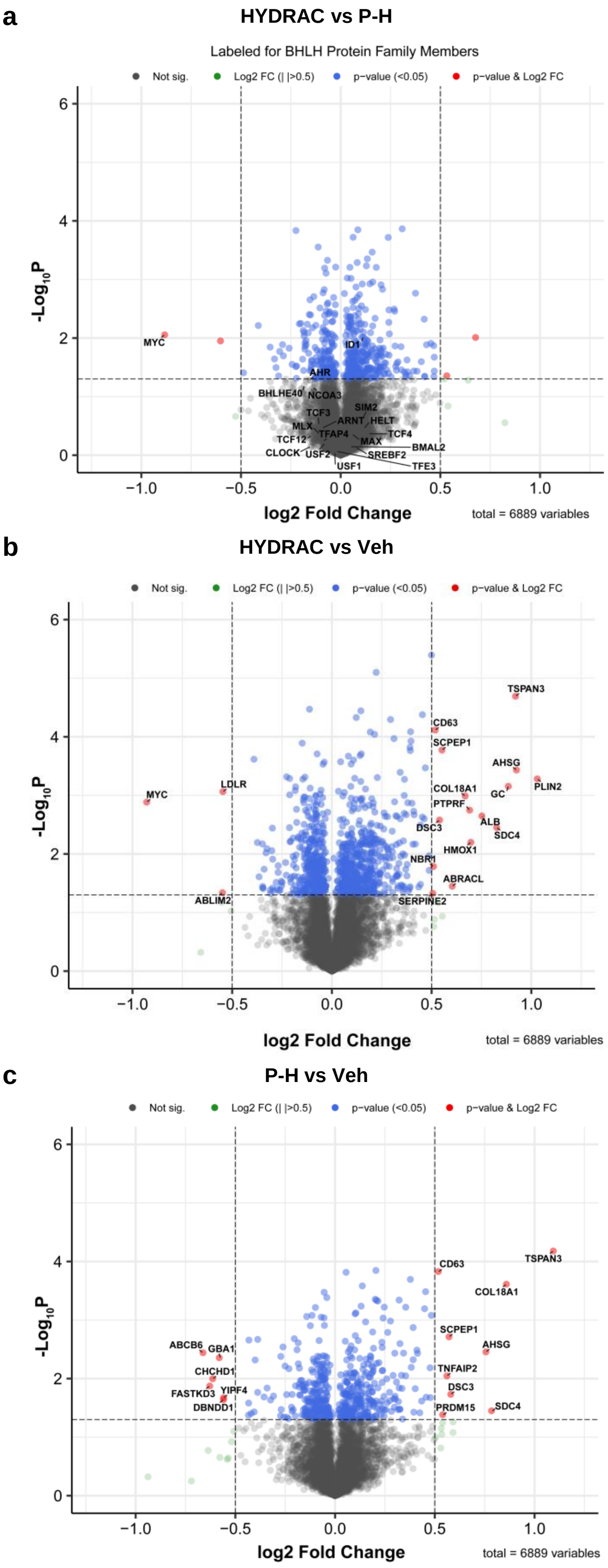


**Supplementary Fig. 13 | TMT-based whole proteome quantification.** Comparisons of PC3 cells treated with HYDRACs vs P-H (**a**), HYDRAC vs Vehicle (**b**), or P-H vs Vehicle (**c**) with proteins showing significant up or down regulation highlighted. BHLH protein family members labeled in (**a**).

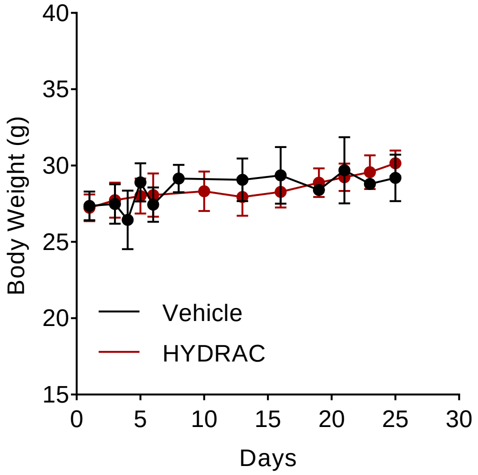


**Supplementary Fig. 14 | Body weights of mice treated with HYDRAC or vehicle.** Mice bearing MycCap allografts were treated with HYDRACs or vehicle (25 mg/kg) 3 times a week and body weight measured over 25 days. N = 5 mice/group.

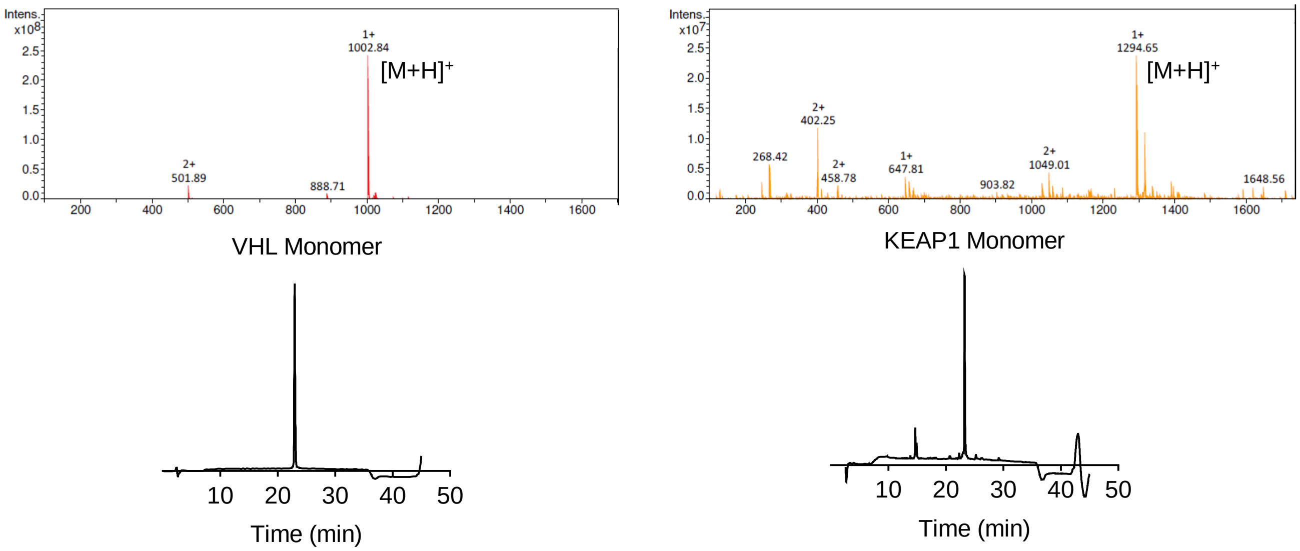


**Supplementary Fig. 15 | Characterization of VHL and KEAP norbornene monomers.** VHL monomer ESI-MS expected $[M+H]^+$: 1002.2, found: 1002.8. KEAP1 monomer ESI-MS expected $[M+H]^+$: 1294.3, found: 1294.6. Analytical HPLC of purified monomers at a 15-65% ACN over 30 min gradient.

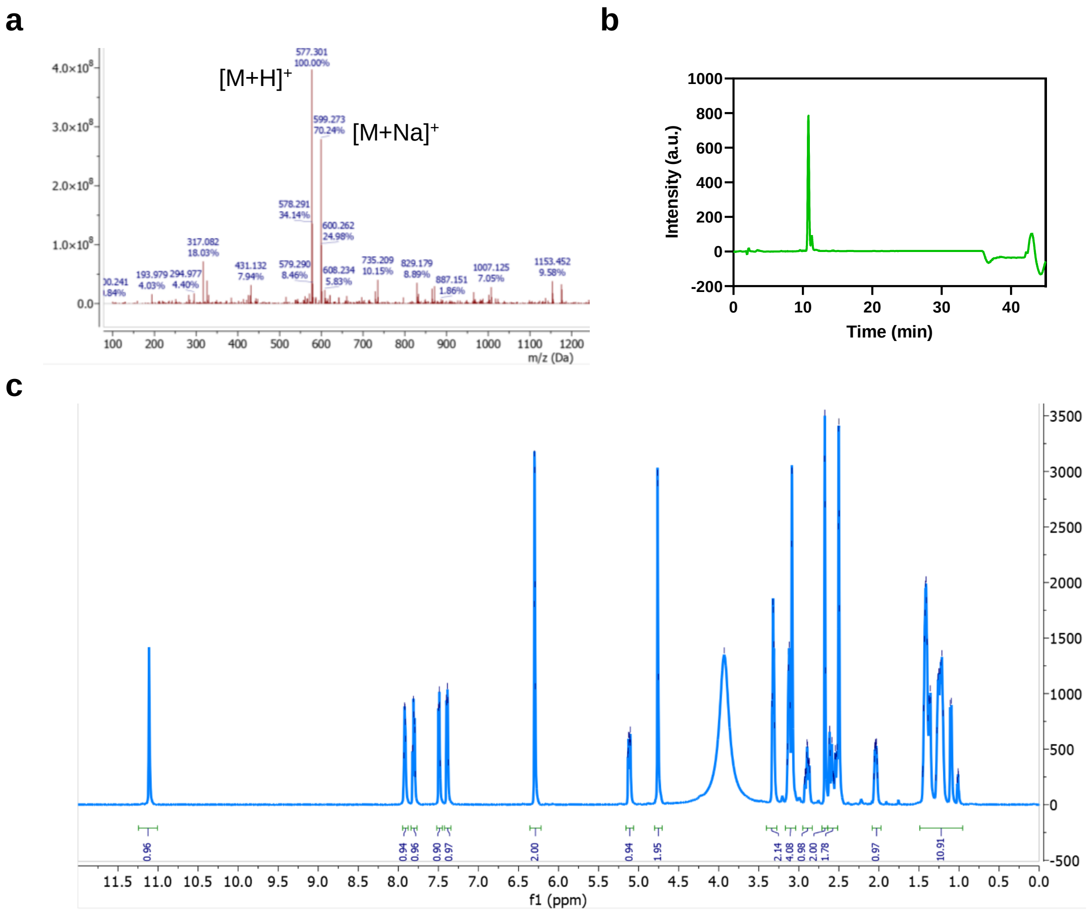


**Supplementary Fig. 16 | Characterization of thalidomide norbornene monomers. a,** ESI-MS expected $[M+H]^+$: 576.22, found 577.3 and $[M+Na]^+$: 599.27. **b,** Analytical HPLC trace with gradient of 35 to 55% acetonitrile. **c,** $^1$H NMR of purified product and peak integrations (See Supplemental Methods).

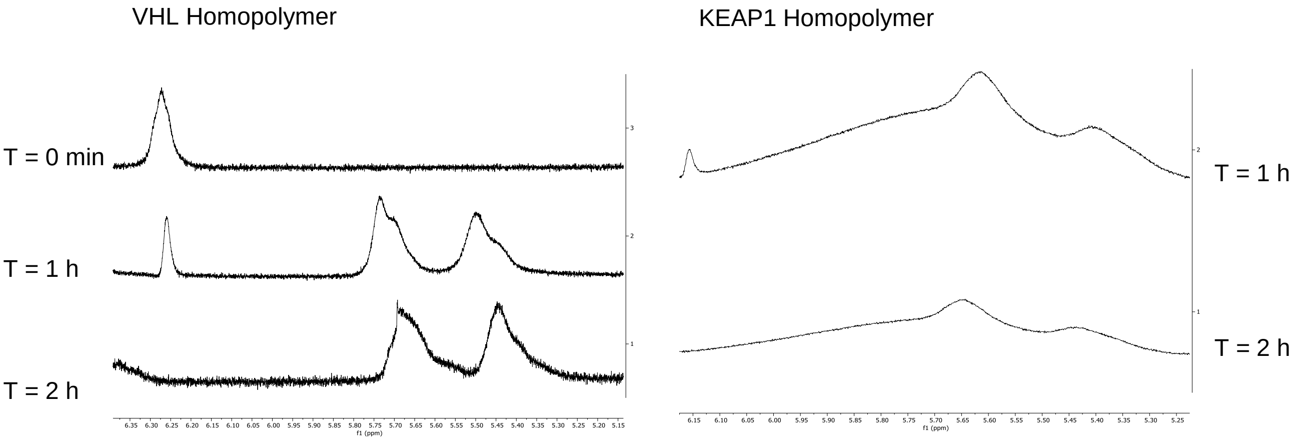


**Supplementary Fig. 17 | Representative $^1$H NMR spectra of VHL and KEAP1 homopolymers.** Resonance at ~δ 6.3 ppm corresponds to monomer olefin protons. Final spectrum recorded at the end of polymerization shows consumption of the monomer. Resonances at ~ δ 5.6 ppm correspond to cis/trans olefinic protons of the polymerized material. Note: Thalidomide homopolymers polymerized near instantly and therefore intermediate spectra could not be obtained.

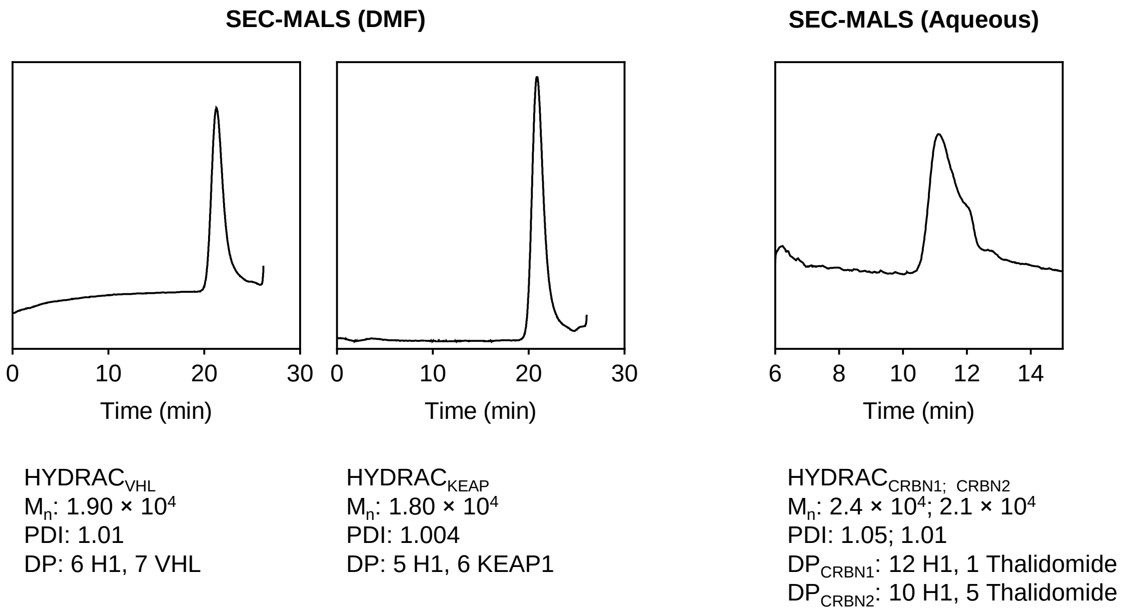


**Supplementary Fig. 18 | Determination of polymer molecular weights by SEC-MALS.** Representative differential refractive index (dRI) chromatograms of HYDRACs containing different E3 ligase recruiters shown.

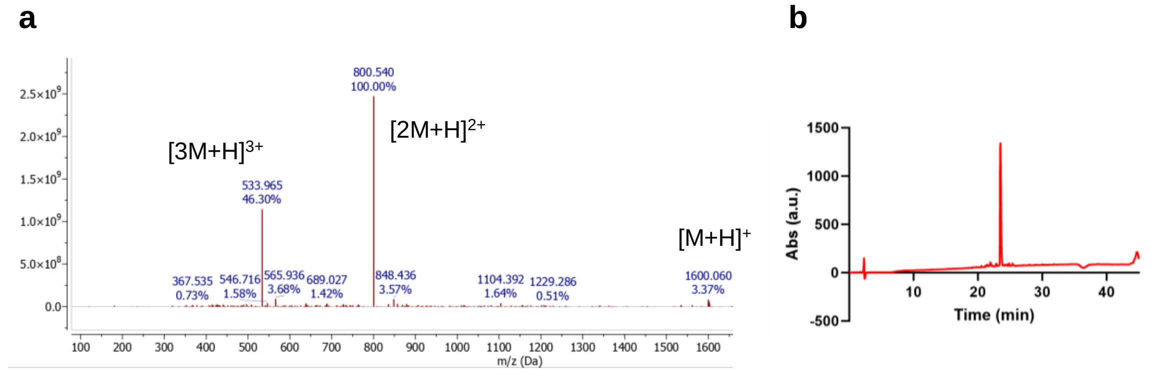


**Supplementary Fig. 19 | Characterization of RAS-targeting norbornene monomers. a,** ESI-MS expected $[M+H]^+$: 1601.6, found 1600.1 and $[M+Na]^+$: 599.27. **b,** Analytical HPLC trace with gradient of 35 to 55% acetonitrile.

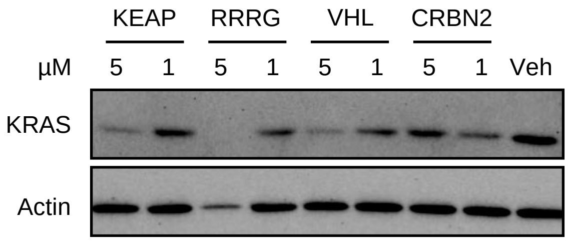


**Supplementary Fig. 20 | Western blot of KRAS protein levels in PANC-1 cells.** PANC-1 cells were treated with RAS-targeted HYDRACs of indicated formulations for 24 h and KRAS protein levels tracked by Western Blot.

| Name | Monomer/Peptide | Mass Calculated | Mass Obtained | Charge at pH = 7 |
|---|---|---|---|---|
| H1 peptide | NELKRAFAALRDQI | 1644.9 | 1645.5 | +1 |
| Nor-H1 | NorAha-NELKRAFAALRDQI | 1904.2 | 1905.2 | +1 |
| Nor-sH1 | NorAha-RQRAIDLFKANELA | 1904.2 | 1904.8 | +1 |
| Nor-RRRG | NorAha-RRRG | 802.9 | 803.2 | +3 |
| Nor-VHL | NorAha-ALAPYIP | 1002.2 | 1002.8 | 0 |
| Nor-KEAP | NorAha-LDPETGEYL | 1294.3 | 1294.6 | -3 |
| Nor-Thalidomide | NorAha-Thalidomide | 576.2 | 577.3 | 0 |
| Nor-RAS | NorAha-FARKTFLKLAF | 1601.6 | 1600.1 | +3 |

**Supplementary Table 1 | Calculated and obtained molecular weight values of peptide monomers used.** Masses obtained via ESI (+) MS. NorAha = N-(hexanoic acid)-cis-5-norbornene-exo-dicarboximide. NELKRAFAALRDQI: H1, ALAPYIP: VHL, LDPETGEYL: KEAP.

| Formulation | Theoretical $M_n$ | $M_n$ | $M_w/M_n$ | Peptide DP |
|---|---|---|---|---|
| P-H | $1.10 \times 10^4$ | $1.2 \pm 0.1 \times 10^4$ | $1.02 \pm 0.02$ | $H1_{5\text{-}7}$ |
| $P\text{-}H_{long}$ | $1.9 \times 10^4$ | $2.3 \times 10^4$ | 1.04 | $H1_{12}$ |
| P-R | $0.5 \times 10^4$ | $0.9 \pm 0.2 \times 10^4$ | $1.01 \pm 0.01$ | $RRRG_{7\text{-}11}$ |
| HYDRAC | $1.90 \times 10^4$ | $2.0 \pm 0.1 \times 10^4$ | $1.01 \pm 0.02$ | $H1_{6\text{-}8}$-*stat*-$RRRG_{6\text{-}10}$ |
| P-sHR | $1.90 \times 10^4$ | $1.9 \pm 0.1 \times 10^4$ | $1.04 \pm 0.02$ | $sH1_{4\text{-}6}$-*stat*-$RRRG_{6\text{-}8}$ |
| $H_1R_9$ | $0.91 \times 10^4$ | $1.1 \times 10^4$ | 1.01 | $H1_1$-*stat*-$RRRG_{10}$ |
| $H_9R_1$ | $1.80 \times 10^4$ | $2.0 \times 10^4$ | 1.001 | $H1_{10}$-*stat*-$RRRG_1$ |
| $HYDRAC_{VHL}$ | $1.75 \times 10^4$ | $1.7 \pm 0.4 \times 10^4$ | $1.01 \pm 0.01$ | $H1_{4\text{-}6}$-*stat*-$VHL_{4\text{-}6}$ |
| $HYDRAC_{KEAP}$ | $1.91 \times 10^4$ | $1.9 \pm 0.1 \times 10^4$ | $1.01 \pm 0.01$ | $H1_{5\text{-}6}$-*stat*-$KEAP_{5\text{-}6}$ |
| $HYDRAC_{CRBN1}$ | $2.35 \times 10^4$ | $2.4 \times 10^4$ | 1.003 | $H1_{12}$-*stat*-$Thal_2$ |
| $HYDRAC_{CRBN2}$ | $2.20 \times 10^4$ | $2.1 \times 10^4$ | 1.05 | $H1_9$-*stat*-$Thal_6$ |

**Supplementary Table 2 | Summary characterization of polymer formulations used.** Tabulated values for the preparation of homopolymers and copolymers. Experimental $M_n$ values obtained by SEC-MALS are listed and align with expected values calculated from targeted DPs. A minimum of three separate synthetic batches were quantified for H, R, HR, and sHR. DPs within 25% of desired value were considered equivalent formulations. Data depict mean ± s.d. Value ranges listed in parentheses. $[M]_0:[I]_0$: initial monomer-to-catalyst ratio used (target DP), $M_n$: number-average molecular weight from light scattering, PDI ($M_w/M_n$): polydispersity index, DP: experimentally determined degree of polymerization.

| Fig | Label | $M_n$ | Composition |
|---|---|---|---|
| 1c,d | P-H | $1.25 \times 10^4$ | $H1_6$ |
| | P-$H_{long}$ | $2.31 \times 10^4$ | $H1_{12}$ |
| | HYDRAC | $2.20 \times 10^4$ | $H1_7$-*stat*-$RRRG_7$ |
| 1e | HYDRAC | $1.75 \times 10^4$ | $H1_6$-*stat*-$RRRG_7$ |
| 1f | HYDRAC | $1.75 \times 10^4$ | $H1_6$-*stat*-$RRRG_7$ |
| 1g | HYDRAC | $1.75 \times 10^4$ | $H1_6$-*stat*-$RRRG_7$ |
| 1h | P-$H_B$ | $1.20 \times 10^4$ | $H1_7$-biotin |
| | $HYDRAC_B$ | $1.80 \times 10^4$ | $H1_7$-*stat*-$RRRG_6$-biotin |
| | P-$sH_B$ | $1.10 \times 10^4$ | $sH1_5$-biotin |
| | P-$sHR_B$ | $1.80 \times 10^4$ | $sH1_7$-*stat*-$RRRG_6$-biotin |
| 2b | P-$H_F$ | $1.06 \times 10^4$ | $H1_5$-$Cy5.5_1$ |
| | P-$R_F$ | $8.60 \times 10^3$ | $RRRG_9$-$Cy5.5_1$ |
| | $HYDRAC_F$ | $1.70 \times 10^4$ | $H1_6$-*stat*-$RRRG_6$-$Cy5.5_1$ |
| | P-$sHR_F$ | $1.60 \times 10^4$ | $sH1_6$-*stat*-$RRRG_5$-$Cy5.5_1$ |
| 2c | $HYDRAC_F$ | $1.70 \times 10^4$ | $H1_6$-*stat*-$RRRG_6$-$Cy5.5_1$ |
| 2d | $HYDRAC_F$ | $1.70 \times 10^4$ | $H1_6$-*stat*-$RRRG_6$-$Cy5.5_1$ |
| 2e | HYDRAC | $1.90 \times 10^4$ | $H1_7$-*stat*-$RRRG_7$ |
| 2f | HYDRAC | $1.90 \times 10^4$ | $H1_7$-*stat*-$RRRG_7$ |
| 3a | HYDRAC | $1.90 \times 10^4$ | $H1_7$-*stat*-$RRRG_7$ |
| | P-sHR | $1.60 \times 10^4$ | $H1_6$-*stat*-$RRRG_5$ |
| 3b | HYDRAC | $1.90 \times 10^4$ | $H1_7$-*stat*-$RRRG_7$ |
| | P-sHR | $1.70 \times 10^4$ | $sH1_6$-*stat*-$RRRG_7$ |
| 3c | HYDRAC | $1.90 \times 10^4$ | $H1_7$-*stat*-$RRRG_7$ |
| | P-sHR | $1.70 \times 10^4$ | $sH1_6$-*stat*-$RRRG_7$ |
| 3d | P-H | $1.00 \times 10^4$ – $1.30 \times 10^4$ | $H1_{5\text{-}7}$ (range) |
| | P-R | $7.00 \times 10^3$ – $1.00 \times 10^4$ | $RRRG_{9\text{-}12}$ |
| | HYDRAC | $1.60 \times 10^4$ – $2.10 \times 10^4$ | $H1_{5\text{-}7}$-*stat*-$RRRG_{6\text{-}9}$ |
| | P-sH | $0.80 \times 10^4$ – $1.20 \times 10^4$ | $sH1_{4\text{-}6}$ |
| | P-sHR | $1.50 \times 10^4$ – $1.80 \times 10^4$ | $H1_{5\text{-}6}$-*stat*-$RRRG_{6\text{-}9}$ |

| Fig | Label | $M_n$ | Composition |
|---|---|---|---|
| 3e | HYDRAC | $1.65 \times 10^4$ | $H1_6$-*stat*-$RRRG_7$ |
| | P-sHR | $1.80 \times 10^4$ | $sH1_5$-*stat*-$RRRG_9$ |
| 3f | $H_9R_1$ | $2.00 \times 10^4$ | $H1_{10}$-*stat*-$RRRG_1$ |
| | $H_1R_9$ | $1.05 \times 10^4$ | $H1_1$-*stat*-$RRRG_{10}$ |
| | HYDRAC | $2.10 \times 10^4$ | $H1_7$-*stat*-$RRRG_8$ |
| 3g | HYDRAC | $1.75 \times 10^4$ | $H1_6$-*stat*-$RRRG_7$ |
| 3h | P-H | $1.10 \times 10^4$ | $H1_6$ |
| | P-sHR | $1.70 \times 10^4$ | $sH1_6$-*stat*-$RRRG_6$ |
| | HYDRAC | $1.80 \times 10^4$ | $sH1_6$-*stat*-$RRRG_8$ |
| 4a | P-H | $1.29 \times 10^4$ | $H1_7$ |
| | HYDRAC | $1.65 \times 10^4$ | $H1_6$-*stat*-$RRRG_7$ |
| | P-sHR | $1.80 \times 10^4$ | $sH1_5$-*stat*-$RRRG_9$ |
| 4b | P-H | $1.15 \times 10^4$ | $H1_6$ |
| | P-R | $0.90 \times 10^4$ | $RRRG_{10}$ |
| | HYDRAC | $1.90 \times 10^4$ | $H1_5$-*stat*-$RRRG_6$ |
| | P-sHR | $1.50 \times 10^4$ | $sH1_6$-*stat*-$RRRG_5$ |
| 4c | HYDRAC | $2.00 \times 10^4$ | $H1_7$-*stat*-$RRRG_7$ |
| 4d | HYDRAC | $1.90 \times 10^4$ | $H1_6$-*stat*-$RRRG_7$ |
| 4e | HYDRAC | $2.00 \times 10^4$ | $H1_7$-*stat*-$RRRG_7$ |
| 4f | HYDRAC | $1.75 \times 10^4$ | $H1_6$-*stat*-$RRRG_6$ |
| 4g,h | HYDRAC | $1.90 \times 10^4$ | $H1_6$-*stat*-$RRRG_7$ |
| 5a,b | HYDRAC | $1.90 \times 10^4$ | $H1_6$-*stat*-$RRRG_7$ |
| 5c | HYDRAC | $1.80 \times 10^4$ | $H1_6$-*stat*-$RRRG_7$-$Cy5.5_1$ |
| 6 | $HYDRAC_{VHL}$ | $1.90 \times 10^4$ | $H1_6$-*stat*-$VHL_7$ |
| | $HYDRAC_{KEAP}$ | $1.80 \times 10^4$ | $H1_5$-*stat*-$KEAP_6$ |
| | $HYDRAC_{CRBN1}$ | $2.40 \times 10^4$ | $H1_{12}$-*stat*-$Thalidomide_1$ |
| | $HYDRAC_{CRBN2}$ | $2.10 \times 10^4$ | $H1_{10}$-*stat*-$Thalidomide_5$ |
| 7 | RAS Homo | $1.10 \times 10^4$ | $RAS_7$ |
| | ${}_{RAS}HYDRAC_{RRRG}$ | $1.80 \times 10^4$ | $RAS_7$-*stat*-$RRRG_8$ |
| | ${}_{RAS}HYDRAC_{VHL}$ | $1.90 \times 10^4$ | $RAS_7$-*stat*-$VHL_8$ |
| | ${}_{RAS}HYDRAC_{KEAP}$ | $2.0 \times 10^4$ | $RAS_7$-*stat*-$VHL_7$ |
| | ${}_{RAS}HYDRAC_{CRBN1}$ | $2.20 \times 10^4$ | $RAS_{14}$-*stat*-$Thal_1$ |
| | ${}_{RAS}HYDRAC_{CRBN2}$ | $1.90 \times 10^4$ | $RAS_{10}$-*stat*-$Thal_5$ |

**Supplementary Table 3 | Experimentally determined molecular weights for polymer batches used in each experiment.** Subscript depicts degree of polymerization for each component piece. *Stat*: statistical copolymer.

### REFERENCES


1. Thompson, M.P. et al. Labelling Polymers and Micellar Nanoparticles via Initiation, Propagation and Termination with ROMP. *Polym Chem* **5**, 1954-1964 (2014).
2. Kammeyer, J.K., Blum, A.P., Adamiak, L., Hahn, M.E. & Gianneschi, N.C. Polymerization of Protecting-Group-Free Peptides via ROMP. *Polym Chem* **41**, 3929-3933 (2013).
3. Blum, A.P. et al. Peptides Displayed as High Density Brush Polymers Resist Proteolysis and Retain Bioactivity. *Journal of the American Chemical Society* **136**, 15422-15437 (2014).
4. Ungerleider, J.L., Kammeyer, J.K., Braden, R.L., Christman, K.L. & Gianneschi, N.C. Enzyme-Targeted Nanoparticles for Delivery to Ischemic Skeletal Muscle. *Polym Chem* **8**, 5212-5219 (2017).
5. Matson, J.B. & Grubbs, R.H. Monotelechelic Poly(oxa)norbornenes by Ring-Opening Metathesis Polymerization using Direct End-Capping and Cross Metathesis. *Macromolecules* **43**, 213-221 (2010).
